# Pre-trauma memory contextualization as predictor for PTSD-like behavior in male rats

**DOI:** 10.1101/2021.07.17.452780

**Authors:** Milou S.C. Sep, R. Angela. Sarabdjitsingh, Elbert Geuze, Marian Joёls

**Author notes:** Corresponding author: Milou S.C. Sep; @; Universiteitsweg 100, 3584 CG Utrecht, the Netherlands.

## Abstract

While many people experience potentially threatening events during their life, only a minority develops posttraumatic stress disorder (PTSD). The identification of individuals at risk among those exposed to trauma is crucial for PTSD prevention in the future. Since re-experiencing trauma elements outside of the original trauma-context is a core feature of PTSD, we investigate if the ability to bind memories to their original encoding context (i.e. memory contextualisation) predicts PTSD vulnerability. We hypothesize that pre-trauma neutral memory contextualization (under stress) negatively relates to PTSD-like behavior, in a prospective design using the cut-off behavioral criteria rat model for PTSD. 72 male Sprague Dawley rats were divided in two experimental groups to assess the predictive value of 1) memory contextualization without acute stress (NS-group) and 2) memory contextualization during the recovery phase of the acute stress-response (S-group) for susceptibility to PTSD-like behavior. A powerful extension to regression analysis-path analysis-was used to test this specific hypothesis, together with secondary research questions. Following traumatic predator scent stress, 19.4% of the rats displayed PTSD-like behavior. Results showed a negative relation between pre-trauma memory contextualization and PTSD-like behavior, but only in the NS-group. Pre-trauma memory contextualization was positively related to fear association in the trauma environment, again only in the NS group. If the predictive value of pre-trauma contextualization of neutral information under non-stressful conditions for PTSD susceptibility is replicated in prospective studies in humans, this factor would supplement already known vulnerability factors for PTSD and improve the identification of individuals at risk among the trauma exposed, especially those at high trauma risk such as soldiers deployed on a mission.

## 1. Introduction

Posttraumatic stress disorder (PTSD) can arise in the aftermath of an intense traumatic experience (American Psychiatric Association, 2013). The intrusive recollections, hyperarousal and avoidance behaviors associated with PTSD (American Psychiatric Association, 2013) put a major burden on the individual and society (Bomyea et al., 2012; Kliem and Kroger, 2013). Interestingly, only a minority of trauma-exposed individuals develop full-blown PTSD. For instance, although over 80 % of the Dutch population encounters a potentially traumatic event during their life, PTSD only occurs in 7.4% of the general population (de Vries and Olff, 2009). This points towards substantial individual differences in the reaction to traumatic experiences and disease susceptibility (Bomyea et al., 2012; Cathomas et al., 2019; Holmes and Singewald, 2013; Yehuda and LeDoux, 2007). Several vulnerability factors prior to trauma have been identified over the past decades that hold potential to identify individuals at risk (Bomyea et al., 2012; DiGangi et al., 2013; Tortella-Feliu et al., 2019; Yehuda and LeDoux, 2007). In the cognitive domain, these factors include low intelligence, poor neuropsychological performance and negative cognitive biases (Bomyea et al., 2012; Kremen et al., 2012). Among genetic and physiological risk factors several are related to HPA-axis functioning (Sheerin et al., 2020; van Zuiden et al., 2013), such as low baseline cortisol, and enhanced negative feedback of the glucocorticoid system via glucocorticoid receptors (GRs) (Fischer et al., 2021; Steudte-Schmiedgen et al., 2015; Szeszko et al., 2018; Turner et al., 2020). Since risk factors, within the individual, cluster around blunted HPA-axis response and decreased cognitive abilities (Bomyea et al., 2012), the interplay between stress and cognition may provide valuable insights. Besides, these systems and the way they interact demonstrate substantial individual differences (Sweis et al., 2013), which makes them suitable to discriminate between individuals.

In this study we used a rat model (Cohen et al., 2012) to further explore PTSD vulnerability factors related to the HPA-axis and cognition. The selected model uses cut-off behavioural criteria for PTSD based on the clinical symptoms of PTSD; and differentiates between resilient and susceptible animals among the trauma-exposed. Using an animal model for PTSD (Verbitsky et al., 2020) has two major advantages over a prospective human cohort study. First, it provides us with the opportunity to induce trauma in a controlled manner, allowing to investigate *differences* in individual response after the *same* traumatic effect. Second, it enables disentangling of vulnerability factors from trauma-induced or PTSD-related behavioural responses, as *pre-* and *post*-trauma behaviours at subject level can be compared.

In our model we focused on one particular cognitive ability, i.e. memory contextualization which is the ability to store declarative memories with details from the circumstances or the environment that surrounded the event. One of the core feature of PTSD, i.e. reexperiencing elements from trauma outside of the original context, seems to indicate that contextualization is impaired in patients with PTSD (Liberzon and Abelson, 2016), which may represent a predisposing trait rather than a state. Direct evidence is lacking (Maren et al., 2013), but PTSD patients indeed show poorer performance on processing of context cues (Kremen et al., 2012). Moreover, hippocampal function is critically involved in the development of context memory (Acheson et al., 2012; Maren et al., 2013; Mitsushima et al., 2011; Rugg et al., 2012) and neuroimaging studies reported hippocampal abnormalities in PTSD patients, even prior to the trauma (Acheson et al., 2012; Gilbertson et al., 2002; Matar et al., 2013). Interestingly, HPA-axis related hormones in humans seem to influence memory contextualization, as exogenously administered cortisol affects the ability to contextualize memories in a time-dependent manner (van Ast et al., 2013). More specifically, rapid effects of cortisol impaired memory contextualization, while the slow effects of cortisol–relevant for the aftermath of stress-improved performance (van Ast et al., 2013). Recently we replicated this experiment with a psychosocial stressor in healthy male subjects, and found that stress affected memory contextualization of neutral, but not emotional, information in a similar time-dependent manner (Sep et al., 2019b). Taken together, this suggests that insufficient contextualization of neutral trauma-related information, possibly due to a blunted HPA-axis response and hence impaired development of slow corticosteroid actions, may be a predisposing trait for PTSD.

The main objective of the current study was thus to determine if neutral memory contextualization, particularly in the aftermath of stress compared to non-stressed conditions, is a stable pre-trauma trait that predicts vulnerability to PTSD at an individual level in male rats. We furthermore formulated the following sub-objectives: i) to determine if the ability to contextualize neutral memories is a stable trait, which would give a basis for retrospective investigations in human cohorts; ii) to examine if pre-trauma neutral memory contextualization predicts fear association in the trauma-related context; and iii) to investigate whether HPA-axis reactivity during trauma mediates the effect of pre-trauma neutral memory contextualization on PTSD susceptibility.

## 2. Methods

### 2.1 Animals and housing conditions

All animal procedures were approved by the Animal Ethical Committee at Utrecht University, The Netherlands (AVD1150020186349) prior to the start of the study. Every effort was taken to minimize animal suffering and all procedures were conducted in accordance with the FELASA guidelines and the EU directive (2010/63/EU). The outline of this study was preregistered on *preclinicaltrials.eu* (identifier PCTE0000150) and performed according to the ARRIVE guidelines.

Six-weeks old, male, test naïve, Sprague Dawley rats (SD, Charles River Laboratories, Germany) arrived two weeks prior to the start of the experiment in the animal facility. During these two weeks, the animals were allowed to acclimatize to the animal facility and handled twice daily for 2 minutes. The SD rats were group-housed in pairs in Type IV Macrolon cages (480×375×210 cm) with Aspen Woodchips bedding and a wood block and Perspex tube as environmental enrichment. The cages were randomly placed on the housing racks in a vivarium with background radio music, temperature (22 ± 2°C) and humidity (45-64%) control, and a normal 12-h light/dark cycle (lights on: 9 am-9 pm). Standard chow pellets (CRM (E), Special Diet Services, UK) and tap water were provided ad libitum. At the start of behavioral testing, SD rats were 8 weeks old and had an average +/-standard deviation weight of 344 +/-47 g (range: 236-450 g). Throughout all procedures non-experimental disturbances were kept to a minimum to avoid any unwanted stress for the animals.

Test naïve, male Long-Evans rats (LE, in-house breeding surplus, n=11), weighing 544-783g (mean 656 g), served as residents to induce mild social stress in the experimental SD animals (see below). LE rats were housed under the same conditions as the SD rats, but in another vivarium in the animal facility, to avoid olfactory, auditory and visual contact between the SD and LE rats prior to the experiment. LE rats were single-housed 10 days prior to a resident-intruder test (RI) and group-housed 2-3 per Type IV Macrolon cage for the remainder of the experiment. Each LE rat had at least 2.5-weeks of rest between subsequent involvement in RIs.

### 2.2 Experimental design

In agreement with the preregistered design, 72 SD rats were randomly assigned to one of two experimental groups (n=36/group) to assess the predictive value of 1) memory contextualization (MC) without acute stress (NS-group) and 2) MC during the recovery phase of the acute stress-response (S-group) for the development of PTSD-like behavior. Each SD animal was an experimental unit and the group size was based on an a priori power analysis using 1) the previously reported small to medium effect of another hippocampus-dependent learning task on PTSD (Goswami et al., 2012), 2) the sample size of a previous prospective study with the same PTSD model (i.e. the cut-off behavioral criteria (CBC) model) that detected a small to medium effect (Danan et al., 2018), and 3) the anticipated prevalence of PTSD-like behavior in the CBC model (Cohen et al., 2018, 2012; Danan et al., 2018).

The experimental SD rats were tested in 9 batches of 8 animals (4 NS-group and 4 S-group per batch). All procedures were performed between 9:30 am and 13:30 pm (light phase) and all animals were habituated to the behavioral testing rooms 24-hours before each behavioral test. The complete experimental timeline for both groups is depicted in Figure 1.

**Figure 1.**
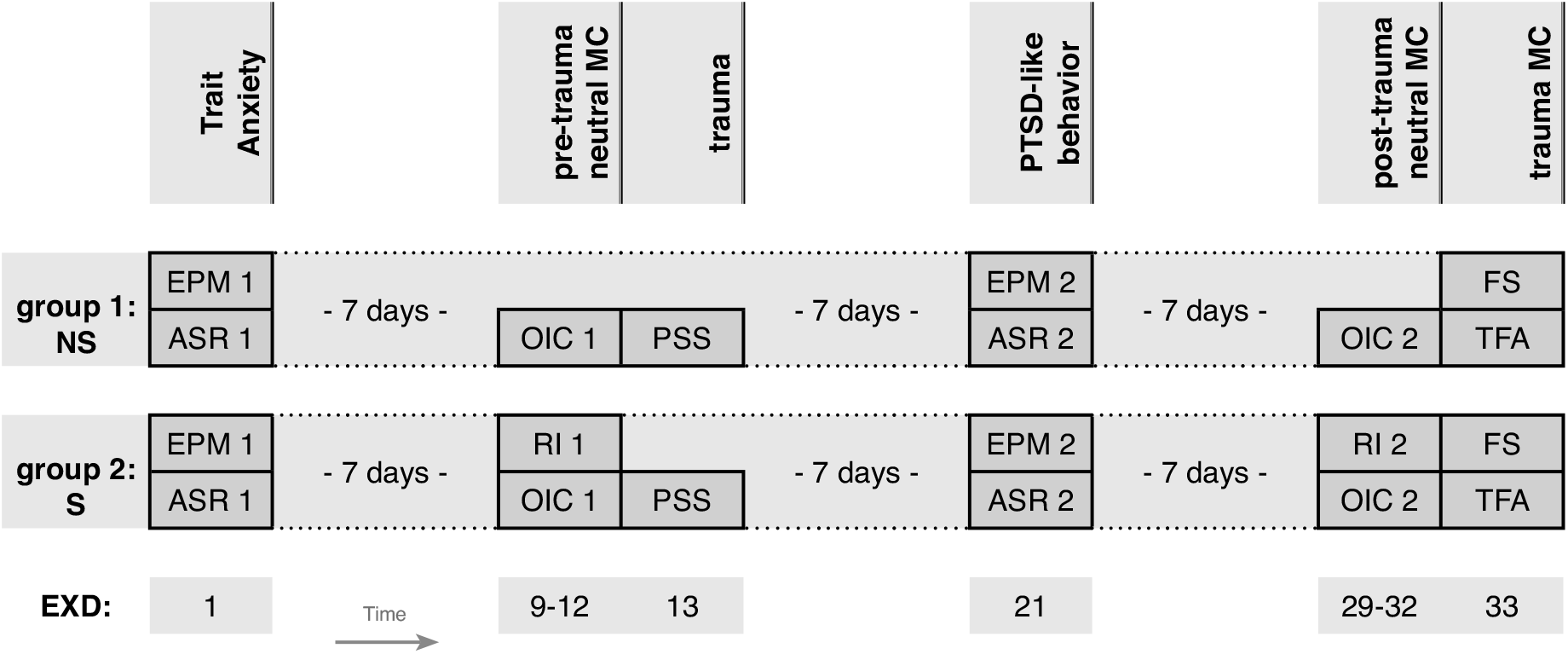
Experimental Timeline per group. ASR: Acoustic Startle Response; EPM: Elevated Plus Maze; EXD: Experimental Day; OIC: Object in Context; NS: no-stress; PSS: Predator scent stress; RI: Resident-intruder; S: stress; TFA: Trauma fear association; FS: fear sensitization. The behavioral tests were conducted in four experimental room: 1) EPM + ASR, 2) RI, 3) OIC, 4) PSS + TG + TCM.

At the start of the experiment (experimental day 1 = EXD 1), pre-trauma basal anxiety (e.g. trait anxiety) was measured in all SD rats. Starting eight days later, pre-trauma neutral MC was assessed over 4 consecutive days (EXD9-12) following learning during either the recovery phase of the stress-response or without mild social stress, in the S- and NS-group respectively. The next day (EXD13) all SD rats were exposed to a traumatic event and after a 7-days’ delay post-trauma anxiety (e.g. PTSD-like behavior) was measured (EXD21). Starting eight days later (EXD29-32), post-trauma neutral MC was measured over 4 consecutive days during the recovery phase of the stress-response (S-group) or without mild social stress (NS-group). The next day (EXD33) trauma MC was measured in all SD rats. Bodyweight was measured before and after each behavioral test (10 timepoints in total). The experimental testing order was randomized per behavioral assessment.

The behavioral tests were conducted in four experimental rooms: anxiety measures in room #1; mild social stress in room #2; neutral MC in room #3; and traumatic event and trauma MC in room #4. Each experimental set-up was built in Duplo (except for the elevated plus maze), which allowed us to prevent additional separation stress by exposing two SD cage mates simultaneously to the experimental procedures. The investigators were not blinded to experimental groups during the experiments, but remained blinded to pre- and post-trauma performance, until all measurements of an individual rat were collected, i.e. data preprocessing and analysis were performed after completion of all experimental procedures. All analyses were performed blindly.

Prior to the start of the experiment, 4 pilots were conducted to validate parts of the experimental set-up with animals of the same strain and age; relevant details are provided in appendix A6 and the text below. In total 42 naïve SD rats and 4 LE rats were used in these pilots (these LE rats were reused in the actual experiment).

### 2.3 Stress Interventions / manipulations

#### 2.3.1 Mild social stress

Rats in the S-group were exposed to mild social stress 150 minutes before neutral MC learning (both before and after trauma), via a mild version of the resident-intruder paradigm (RI) without physical contact (Blanchard et al., 2001). Methodological RI details are provided in appendix A1. This social stressor and timepoint were selected to align with earlier experiments in humans, where the time-dependent effects of social stress on memory contextualization were investigated using the Trier Social Stress Test (Joёls et al., 2018, 2012; Sep et al., 2019a, 2019b). Plasma corticosterone increase from baseline, as proxy for HPA-axis activation, in SD rats by the RI procedure was confirmed in two separate pilot experiments (Appendix 6: Pilot I and IV). Animals in the NS-group stayed in their home cage and were not exposed to the social stress intervention. Compared to the control NS-group, therefore, the S-group were exposed to a combination of the RI-test and removal of their home cage environment for the duration of the intervention.

#### 2.3.2 Traumatic predator scent stress

To model a traumatic experience, all SD rats (S- and NS-group) were exposed to a cat urine predator scent stressor (PSS) for 10 min (for details see appendix A2). It has been shown that this model elicits the PTSD phenotype in cat-naïve rats (e.g. refs: (Cohen et al., 2018; Danan et al., 2018)). This type of trauma was selected because it represents a ‘natural’ potentially life-threatening situation for the rat (Cohen et al., 2012). A life-threating situation is a key traumatic experience that can induce PTSD in humans, according to the clinical diagnostic criteria (American Psychiatric Association, 2013). Each exposure cage was placed in an opaque box to prevent visual contact between the rats during PSS exposure. Plasma corticosterone increase from baseline in SD rats by PSS exposure was validated in two separate pilot experiments (Appendix 6: Pilot I and IV).

### 2.4 Neuroendocrine measurements: Measurement of plasma corticosterone

To determine HPA-axis reactivity to PSS (and RI, in pilot experiments), small aliquots of blood were collected-via a tail cut (Fluttert et al., 2000)-directly before and 30 minutes after the start of PSS exposure in Microvettes (CB300, Starstedt, Germany) on ice. Although plasma samples are subject to ultradian variations (Lightman et al., 2008), the amplitude of ultradian pulses is very low at the time of day that experiments were performed, so that samples probably quite reliably reflect stress-induced variation. Samples were centrifuged at 5000 rpm for 10 minutes at 4°C. Plasma was isolated and stored at −20°C until corticosterone levels were determined using a commercially available radioimmunoassay (RIA) kit (MP biomedicals, the Netherlands). The difference between corticosterone levels in the two samples per rat was used as an index for HPA reactivity.

### 2.5 Behavioral measurements

#### 2.5.1 Anxiety: EPM & ASR

Exploratory behavior in the elevated plus maze (EPM) and acoustic startle responses (ASR) were measured pre- and post-trauma in all SD animals as an index for anxiety. Appendix A3 and A4 provide methodological details on the EPM and ASR, respectively. Test-retest reliability of these anxiety measures at an 18-day interval was validated in a separate pilot experiment (Appendix A6: Pilot II).

#### 2.5.2 Peri-trauma fear

All movements during PSS exposure were recorded by an overhead video camera, and freezing behavior (the absence of all movement, except breathing (Cohen et al., 2008)) was scored with the activity analysis feature in EthoVision^®^ XT 11.5 (Noldus Information Technology BV, Wageningen, The Netherlands). Less than 0.05% pixel change between consecutive images (i.e. video screenshots) was scored as freezing (Analysis parameters, based on (Pham et al., 2009): sample rate (8.33 samples/sec); detection method (dynamic subtraction; contrast: 19-255); activity (activity threshold: 7; background noise filter: off; compression artifacts filter: on; activity averaging interval: 5). Freezing instances shorter than 0.50 s were excluded. Total cumulative freezing time during exposure to PSS was used to calculate a percentage of total exposure time.

#### 2.5.3 Neutral memory contextualization

SD rats’ ability to contextualize neutral information pre- and post-trauma was measured with two versions of the widely used object-in-context task (OIC) (Dix and Aggleton, 1999): 1) *OlC-round* and 2) *OlC-square*. All animals were exposed to both versions, the order was randomized across animals. Two separate pilot experiments (Appendix 6: Pilot III and IV) were conducted to develop the two versions of the OIC and validate test-retest reliability at an 18-day interval.

#### 2.5.4 Context-dependent trauma memory

Context-dependent memory of the traumatic event was measured 20 days after PSS exposure, by re-exposing the SD rats to the original PSS context (with clean sand, e.g. (Cohen et al., 2018, 2014)) and to a new context with similarities to the PSS context (on the same day). Two new contexts were created by applying vertical lines of black plastic tape to two transparent cages (these transparent cages are identical to the PSS exposure cages (±25×25×40 cm), but never used for PSS exposure). No bedding was provided in the new contexts. First, SD rats were allowed to explore the new context for 5 minutes. After a brief delay in their home cage (for the purpose of cleaning and changing set-up), rats were exposed to the original PSS context (5 minutes). All movements in both contexts were recorded by an overhead video camera, and freezing behavior was quantified as described in section 2.5.2. Percent freezing in novel context was used as index for fear sensitization (FS) and percent freezing in original context as index for trauma fear association (TFA).

### 2.6 Behavioral profiling of PTSD-like behavior

Using the well-validated cut-off behavioral criteria (CBC) model for PTSD (Cohen et al., 2012, 2005, 2004, 2003), post-PSS behavior on the EPM (anxiety index) and ASR (average startle amplitude and percent habituation) served as an index for PTSD-like behavior. This approach was selected because it is based on PTSD symptoms in humans and therefore has high translational value to the clinic. Post-trauma behavior on the EPM and ASR was measured after a 7-days’ delay because 1) one week in a rat-lifetime translates to a month in a human-lifetime (clinical symptoms following a traumatic event need to persist for at least one month to qualify for a PTSD-diagnosis), and 2) earlier studies with the CBC model in rodents observed that behavioral changes at this timepoint are unlikely to change significantly over the next 30 days (Cohen et al., 2012; Richter-Levin et al., 2019).

For statistical analysis, two continuous composite scores were created for baseline pretrauma anxiety (trait anxiety = z(EPM index) + z(ASR Amplitude)) and post-trauma anxiety (PTSD-like behavior = z(EPM index) + z(ASR Amplitude) + z(ASR Habituation)). Note that the ASR Habituation index was not included in the trait anxiety score, as human studies on the relation between trait anxiety and habituation are debated (Campbell et al., 2014; Donaldson, 2014; Grillon et al., 1993; Martin-Soelch et al., 2006).

For descriptive purposes, an adapted version of the CBC-model (Ardi et al., 2016; Horovitz et al., 2014) was used for the behavioral profiling of ‘*affected*’ and ‘*unaffected*’ animals. Animals that showed extreme behavior (defined as beyond cutoff) on three -or more-of the four post-trauma anxiety measures (i.e. EPM open arm entries, EPM time spent in closed arms, ASR amplitude and ASR habituation), were considered *affected*. The cutoff-values for extreme behavior were based on the distribution of pre-trauma performance on the parameter under study: EPM-open entries: below 25th percentile of pre-trauma EPM entries in open arms; EPM-time closed: above 75th percentile of pre-trauma EPM time closed arms; ASR-amplitude: above 75th percentile of pre-trauma ASR amplitude; ASR-habituation: above 75th percentile of pre-trauma ASR habituation (based on a recent observation in humans that PTSD-symptom severity has a positive relation with amygdala habituation to fearful stimuli (Y. J. Kim et al., 2019)).

### 2.7 Statistical analysis / Data analysis

All preprocessing steps and analyses were performed in R version 4.0.3 (R Core Team, 2020), with general aid of packages *tidyr* (Wickham, 2020), *dplyr* (Wickham et al., 2020) and *osfr* (Wolen et al., 2020). Analyses were performed using the *lavaan* package (Rosseel, 2012), and visualizations are created with packages *ggplot* (Wickham, 2016), *gghalves* (Tiedemann, 2020), *RColorBrewer* (Neuwirth, 2014), *waffle* (Rudis and Gandy, 2019), *patchwork* (Pedersen, 2020), *lavaanPlot* (Lishinski, 2018) and *semPlot* (Epskamp, 2019). Data and code are available via Open Science Framework (https://osf.io/v7m28/).

Scores that deviated > 3.29 SDs from the sample mean of a variable were considered outliers and winsorized accordingly (outliers in 6 animals, on 8 different variables) (Tabachnick and Fidell, 2012). Subsequently composite scores were calculated for Trait anxiety (TA) and PTSD-like behavior. Proportion of maximum scaling (POMS) was used to bring the included measures-with exception of the composite scores-to the same metric (Moeller, 2015), i.e. the minimum value was set to 0 and the maximum value set to 1.

The hypothesized direct and indirect relations between the measured variables were combined into a model. Correspondence between this model and the experimental data was tested using path-analysis (α=0.05). This is an extension to multiple regression allowing the examination of more complicated and realistic models (Streiner, 2005) and simultaneous estimation of statistical parameters (path-coefficients) for direct and indirect (i.e. mediation) effects (for examples see (Fujita et al., 2019; Gündner et al., 2019; S. R. Kim et al., 2019)). To build a complete and representative model (Streiner, 2005), specified direct and indirect relations to answer the primary research questions were complemented with 1) relations that result from the experimental design (e.g. batch as covariate), or 2) direct and indirect relations between (other) measured variables that could be expected based on previous studies (e.g., trait anxiety and PTSD). Table A1 (Appendix A8) provides an overview of all included relations (with references where applicable) and the complete hypothesized model is depicted in the path-diagram (Figure 2). Importantly, only the parameters of relations with pre- and post-trauma neutral MC were estimated within each experimental group, as we assumed that only these variables are affected by the experimental difference between groups, i.e. exposure to RI prior to OIC training; all the other estimated parameters were constrained to be equal between the two experimental groups, to limit the number of estimations in the analysis. Note, within-group analysis without constrained parameters yielded similar results (data not shown).

**Figure 2.**
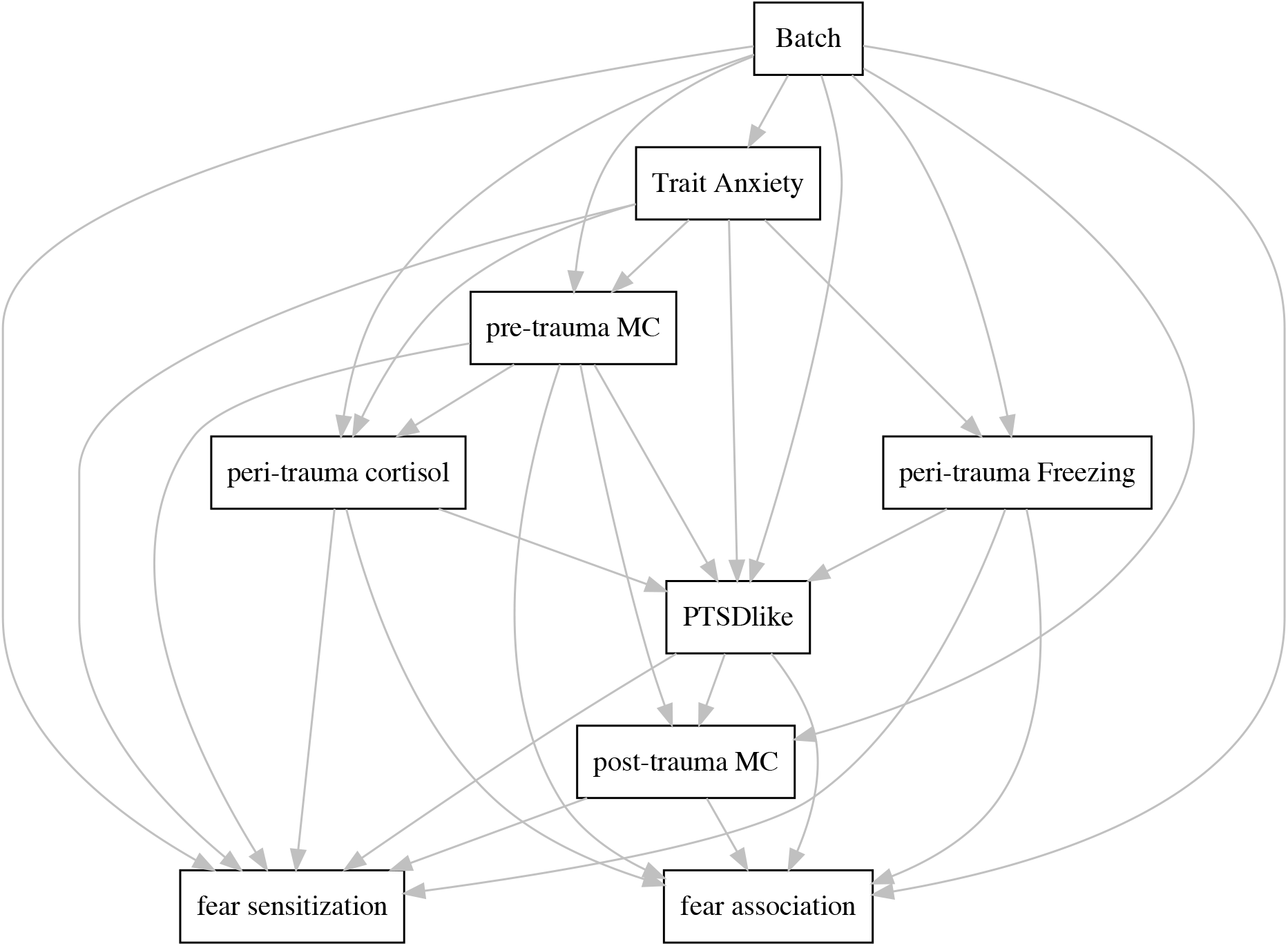
Path-diagram of hypothesized model. Graphical overview of the hypothesized relations that were tested with path-analysis. Relations with pre- and post-trauma MC were estimated within experimental group, all other relations were estimated in the total sample. Note, only regressors are shown in this diagram, not intercepts and covariances.

The assumptions for path-analysis were checked (Streiner, 2005). *Normal distribution* of the variables in the hypothesized model (based on *skewness and kurtosis*) was achieved after square-root-transformations for neutral MC and freezing measures (PSS, TCM, TG) and log-transformation of the shift-transformed (+1) corticosterone difference scores (i.e. *log*(*score+1*)). There was no problematic *multicollinearity* between the variables in the hypothesized model (Maximal *Variance inflation* factor (VIF) was 1.54406 (O’brien, 2007)). Note, data collinearity was tested in a separate main effects regression model.

The model was fitted with Full Information Maximum Likelihood (FIML) estimation in path-analysis, to use all available data (missing values: NS-group = 1, S-group = 3). Bootstrapping (10000 samples) was used to calculate the standard errors and confidence intervals of the direct and indirect paths. Unstandardized (B) and standardized (beta) path-coefficients, indicating relative effects within the model, are reported in the Results. Unstandardized coefficients -and corresponding standard errors and tests- are similar between groups if parameters were constrained to be equal between experimental groups, but standardized coefficients can vary slightly between groups as they are standardized across the complete model for a group.

## 3. Results

### 3.1 Sample characteristics and prevalence of PTSD-like behavior

The animals in the sample varied in bodyweight and behavioral performance, which is essential for the research question at hand: Variation in the no-stress (NS) and stress exposed (S) groups are depicted in Figure 3. According to behavioral profiling following the CBC-model (Figure 4), 14 rats showed PTSD-like behavior (19.4%) following PSS, 58 rats were unaffected (80.6%). Similar percentages were observed after the exclusion of outliers (<3.29 SD): 10 affected animals (15.2%) versus 56 unaffected animals (84.9%). Importantly, these percentages correspond to earlier observations, validating the CBC-model (Cohen et al., 2020, 2018; Danan et al., 2018). For subsequent analyses, all animals were included.

**Figure 3.**
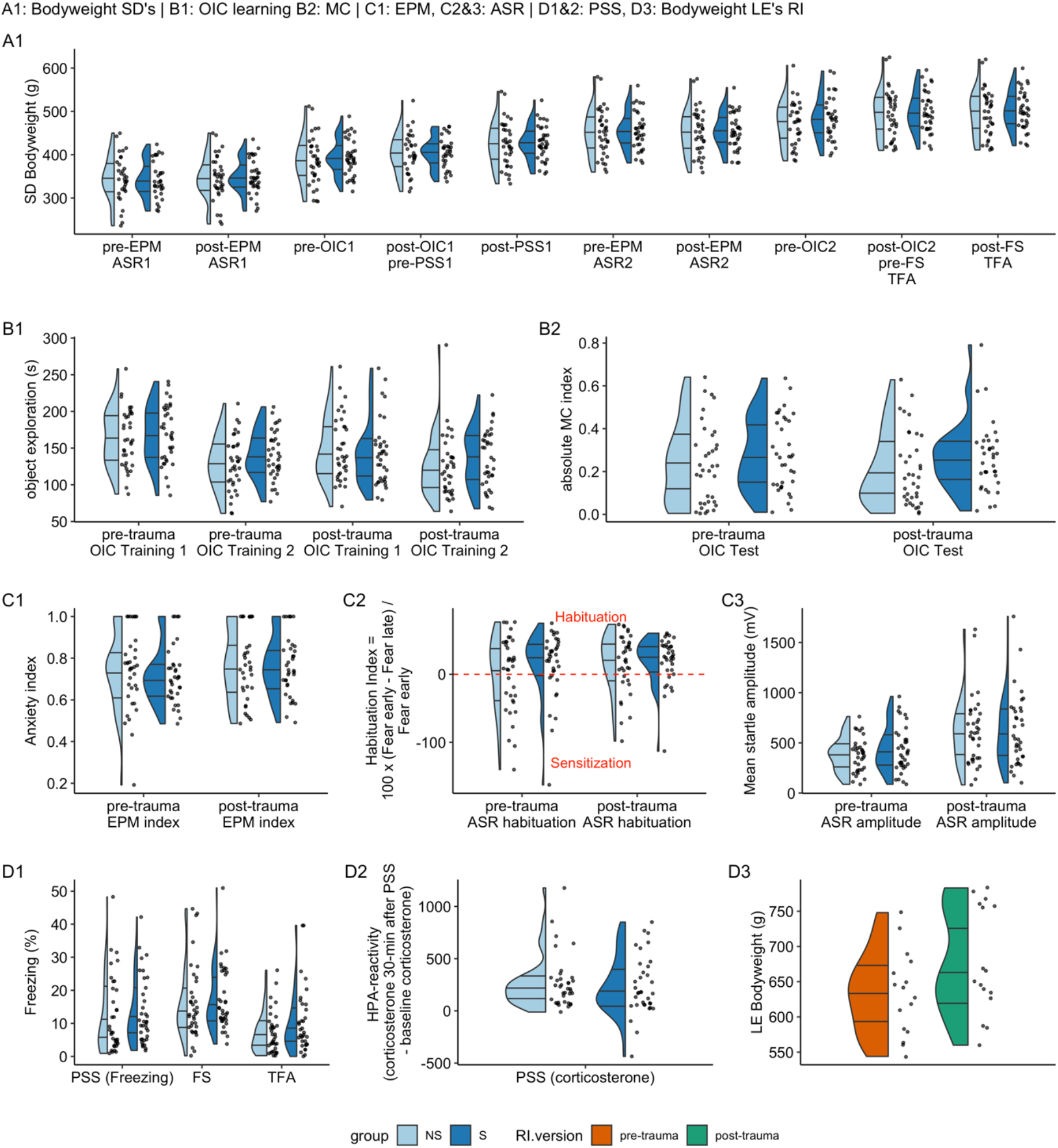
Distribution of bodyweight and behavioral performance: half-violine and half-scatter plots. The bodyweight of Sprague Dawley rats (A1) and Long Evans rats (D3) throughout the experiment and Sprague Dawley rats’ performance on behavioral tasks pre- and post-trauma is shown per experimental group (Object in Context (OIC): B1, B2; Elevated Plus Maze (EPM): C1; Acoustic Startle Response (ASR): C2, C3; Predator scent stress (PSS): D1, D2; fear sensitization (FS): D1; Trauma fear association (TFA): D1). Dots indicate individual animals.

**Figure 4.**
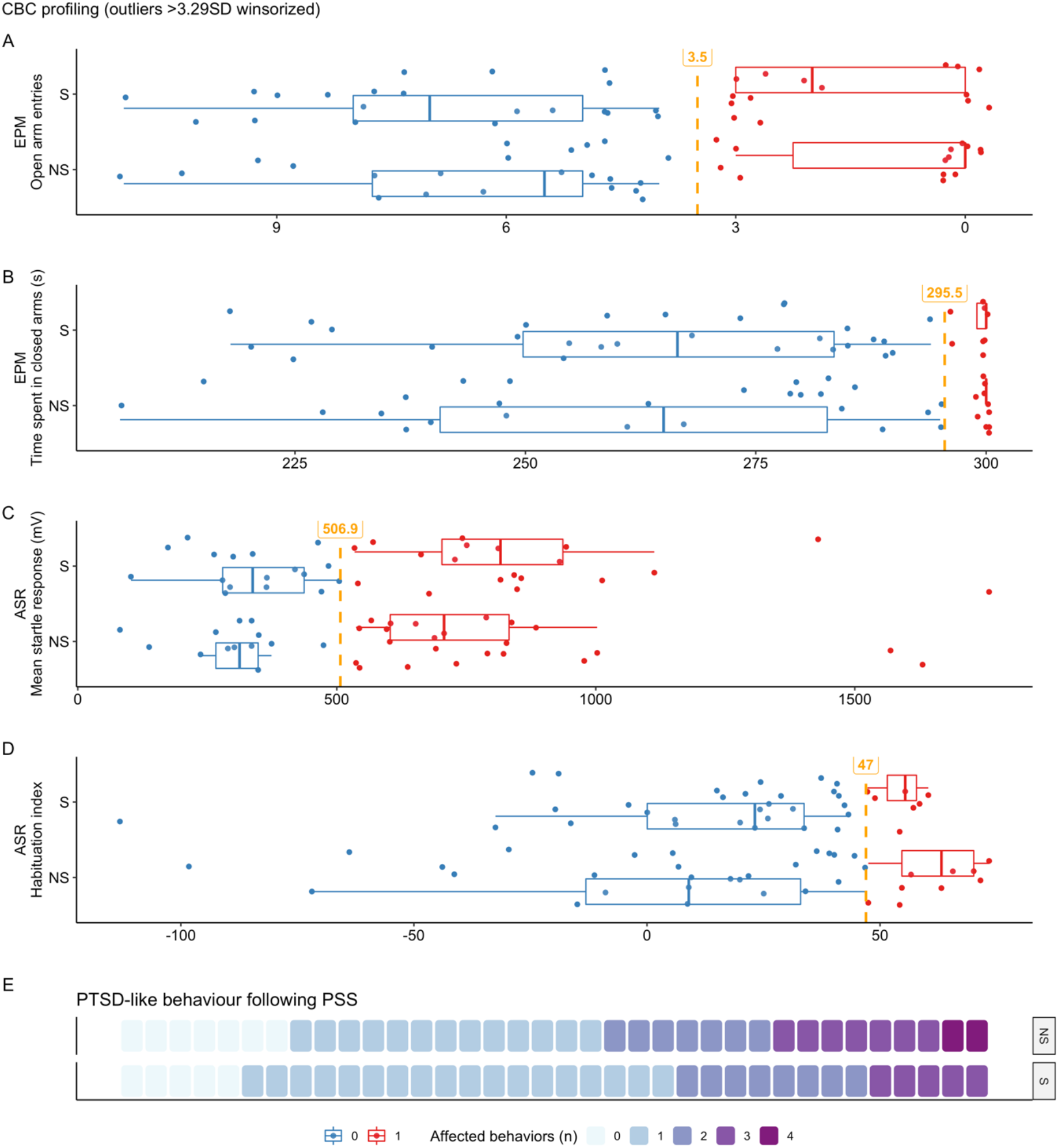
PTSD-like behavior: CBC-Behavioral profiling. Individual performance on the four post-trauma anxiety measures is shown per experimental group (A-D). Values > 3.29SD are winsorized. Dots represent individual animals. The cut-off values for extreme behavior-based on the distribution of pre-trauma behavior- are indicated with dashed yellow lines, extreme behavior is indicated in red and unaffected behavior is shown in blue. Boxplots indicate median with 25^th^ and 75^th^ percentiles (upper and lower hinges). Whiskers indicate 1.5 x inter-quartile range. Per experimental group, the number of affected behaviors per animal in shown in E. PTSD-like behavior ( ≥ 3 affected behaviors) was observed in 9 animals in the NS-group, and in 4 animals in the S-group.

### 3.2 Path-analysis

The full hypothesized model (see Figure 1 and Table B1, Appendix B1) showed a good fit to the experimental data (Hooper et al., 2008), evaluated via goodness-of-fit index (GFI = 0.989), comparative fit index (CFI = 1), Tucker-Lewis Index (TLI = 1.037), root mean square error of approximation (RMSEA = 0; 90%CI = 0 - 0.111), Akaike’s (AIC=323.939) and Schwarz’s Bayesian information criterion (BIC=474.199). No alterations were made to the hypothesized model based on model fit or modification indices. The estimated parameters (including path-coefficients, standard errors, 95% confidence intervals and p-values) for all paths in the hypothesized model (except for covariate ‘batch’) are shown in Table B1 (Appendix B1).

Hypotheses involving pre- and post-trauma neutral memory contextualization (MC) were studied separately for the NS- and S-groups. Our overarching hypothesis was that pretrauma neutral MC negatively relates to PTSD-like symptoms. Path-analyses indeed show that pre-trauma MC had a negative relation with PTSD-like behavior in the NS-group (B = - 1.827, p = .045), though not in the S-group (B = 0.075, p = .936; see Figure 6). As follow-up, Figure 5 shows the correlations between pre-trauma neutral MC and PTSD-like behavior in both groups.

**Figure 5.**
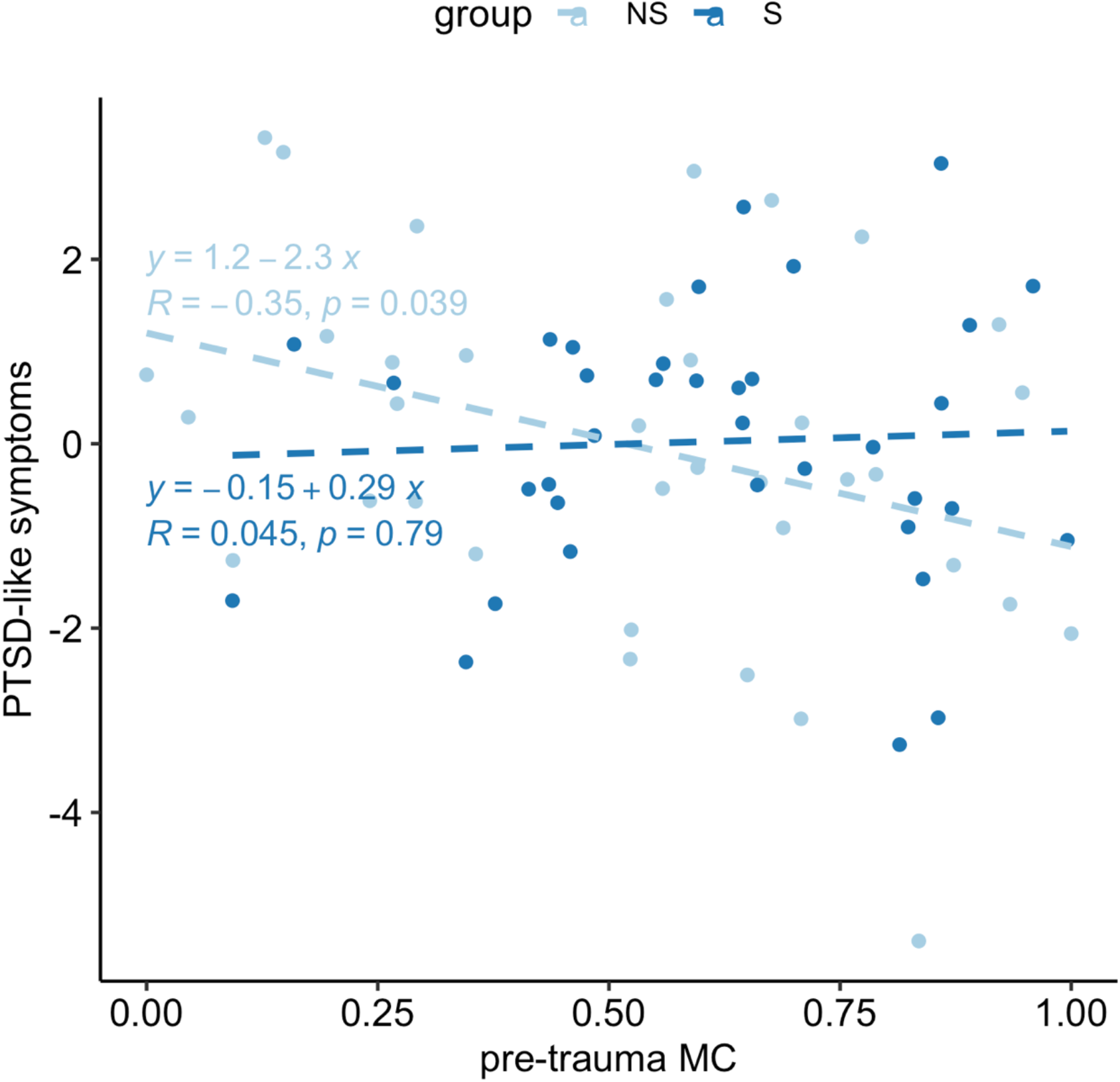
Correlation between pre-trauma neutral MC and PTSD-like symptoms in the NS- and S-groups. We only observed a significant negative correlation in the NS-group, indicating that pretrauma MC measured in the absence of stress had predictive value for PTSD-like behavior, while pretrauma MC determined during the recovery phase of the stress did not.

**Figure 6.**
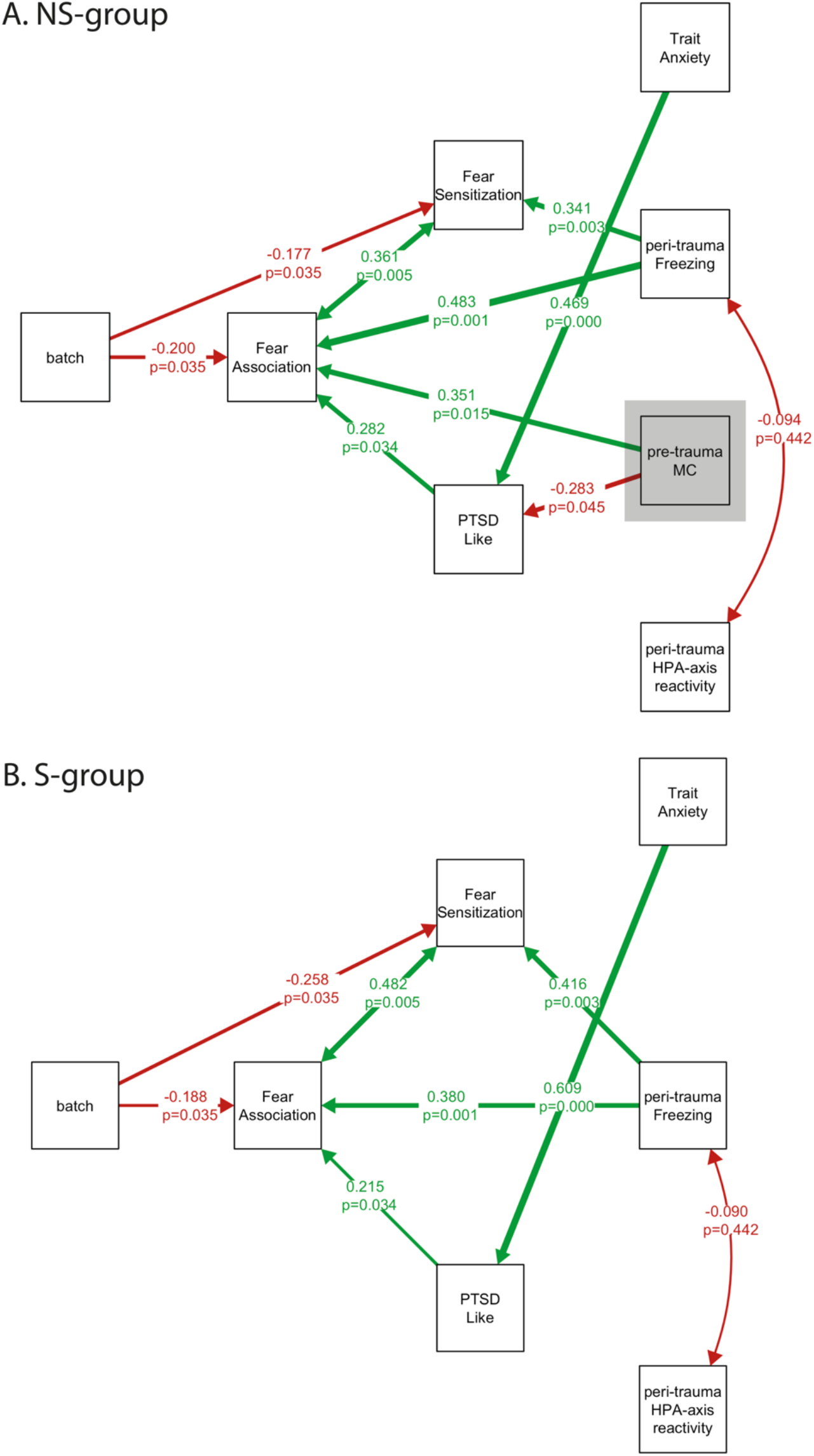
Covariances and significant regression paths with standardized path-coefficients (beta) and p-values per experimental group. Positive relations are visualized with green paths and negative relations with red paths for the NS-group (A) and the S-group (B) separately. In the NS-group (A), pre-trauma MC (highlighted with gray box) predicted PTSD-like behavior (B=-1.827, beta=-0.283, p=0.045) and fear association (B=0.237, beta = 0.351, p=0.015). These relations were not present in the S-group (B). Note, the unstandardized (B) coefficients of the paths that are not connected to MC are equal across groups (as they were constrained within the model). However, the standardized (beta) coefficients-displayed here-can slightly differ between groups because they represent the relative importance of a path within the group. An overview of the unstandardized (B) and standardized (beta) coefficients for all paths-in both groups-is provided in supplementary table 2.

Our first sub-hypothesis was that pre-trauma neutral MC was related to post-trauma neutral MC in both experimental groups. This was not confirmed, neither in the NS-group nor in the S-group (Table B1, Appendix B1). This suggests that neutral MC is not a stable trait, when animals are exposed to a traumatic event. Our second sub-hypothesis assumed that there is a positive relation between pre-trauma MC and fear association (=freezing in the trauma context). This was confirmed in the NS-group (B=0.237, p=.015) but not in the S-group (B=- 0.120, p=.415). Finally, we hypothesized a role of HPA-axis reactivity during trauma, for which we found no significant evidence, neither in the NS-group nor in the S-group.

Apart from MC–which was estimated in a NS group versus the late aftermath of stress on purpose - all other parameters were constrained to be equal between the NS- and S-groups, so that relations were estimated for the complete sample. The analyses resulted in a number of correlations that had not been included in the a priori hypotheses. Firstly, pre-trauma trait anxiety was found to be positively related to PTSD-like behavior (B=0.637, p<0.001). Furthermore, peri trauma freezing showed a positive relation to fear sensitization (B=0.303, p=0.003) and association (B=0.380, p<0.001). Finally, PTSD-like behavior was positively related to fear association (B=0.029, p=0.034). No mediation effects were found (Table B1, Appendix B1). Figure 6 summarizes the main covariances and significant regression paths revealed in the entire path-model, displayed for the NS- and the S-groups separately.

## 4. Discussion

The primary aim of this study was to test if the ability to contextualize neutral information (MC) before trauma predicts susceptibility to PTSD-like behavior after trauma in an animal model. This hypothesis was tested in two experimental groups, where MC was measured 1) during the late aftermath of the stress response (S-group), or 2) without acute stress (NS-group). Testing this specific hypothesis was complemented with secondary research questions in a path-analysis model, a powerful extension to regression analysis. The main finding is that pre-trauma MC indeed has predictive value for PTSD-like behavior, but only in the NS-group. A second finding was that pre-trauma MC correlates positively with fear association in the trauma environment, again only in the NS group. Other secondary hypotheses were rejected.

### Study design

We approached our research question in an animal model for PTSD. This has the advantage (over human experiments) that trauma can be induced in a highly controlled manner, so that all individuals are subjected to the exact same trauma. Moreover, one can expose individual animals to a series of pre-trauma tests, allowing comparison of pre- and post-trauma behaviours at a subject level. We considered this animal study as an exploratory step between the earlier studies on stress and MC in healthy male human subjects (Sep et al., 2019b; van Ast et al., 2013) and future studies in trauma-exposed humans, e.g. soldiers deployed on a mission (van der Wal et al., 2019).

Obviously, any animal model for human psychopathology should be interpreted with great care, since ‘symptoms’ can only be inferred from behavioural expressions. The model we selected (Ardi et al., 2016; Cohen et al., 2012; Horovitz et al., 2014), though, uses cut-off behavioural criteria for PTSD based on the clinical symptoms of PTSD; and differentiates between resilient and susceptible animals among the trauma-exposed. In accordance with the model (Ardi et al., 2016; Cohen et al., 2018; Danan et al., 2018) and the incidence of PTSD in trauma-exposed humans (Kessler et al., 2017; van der Wal et al., 2019), we observed that only 15-20% of the animals met the a priori set criteria for PTSD-like behaviour. This lends credibility to the model and is superior to an often-used approach in which all trauma-exposed animals are compared to non-trauma exposed controls (Richter-Levin et al., 2019).

In our design we followed the earlier human studies (Sep et al., 2019b, 2019a) as closely as possible. For example, we made use of an acute stressor in rats-the resident-intruder situation with a partition between the two rats-which translated the human situation of the Trier Social Stress Test to an ecologically relevant version in rodents, maintaining important elements of social evaluation in the absence of a physical stressor, although there are of course also important differences between the two situations. Noteworthy, the S-group was exposed to a combination of novelty stress and the resident-intruder paradigm, compared to the NS-group which was left undisturbed. Given the somewhat unexpected finding that pretrauma MC was predictive for PTSD susceptibility in the NS-group but not in the S-group, we wondered if the stressor in the latter group had indeed been effective. The observed rise in corticosterone levels post-RI exposure during a pilot study supports the stressful nature of the situation (Figure A5.C, Appendix A6). Moreover, although the study was not designed to examine stress effects on MC–and therefore underpowered to do so-we do have circumstantial evidence that the MC-promoting delayed effects of stress observed in humans (Sep et al., 2019b; van Ast et al., 2013) were also witnessed in rodents. While results from pre-or post-trauma MC separately did not confirm effects of stress exposure, these datasets when combined did confirm a significant positive late effect of stress on MC in the combined sample (Figure B1, Appendix B2). Since pre- and post-trauma MC were found to represent independent variables, one can argue that the combination of datasets is a valid approach. We tentatively conclude that the absence of a predictive value of MC in the S-group therefore is unlikely to be explained by insufficient stress exposure.

In further parallel between the human and rodent setting, we selected a relatively nonstressful rodent memory task that taps into similar hippocampal circuits as the contextual task used in humans (Sep et al., 2021, 2019b). Moreover, similar to research in humans, we determined trait anxiety of the rats, allowing us to correlate this ‘personality trait’ with trauma response and PTSD-susceptibility. Finally, we restricted our study to male rats, since earlier human studies on memory contextualization were also carried out in males (Sep et al., 2019b; van Ast et al., 2013). In this instance this is relevant, since future prospective studies on PTSD in humans seem most feasible in–primarily male-soldiers deployed on mission, given the possibility to extensively test individuals prior to potential trauma exposure (van der Wal et al., 2019). Of note, gender differences in stress-response and in the interaction between stress and cognition are well-documented (Cornelisse et al., 2011; van Ast et al., 2012; Wolf et al., 2001). Moreover, a recent study with male and female rats found that the blunted HPA-axis response to trauma, an important component of our hypothesis, was only present in males (Pooley et al., 2018). All in all, the current design only sheds light on the situation in male subjects and cannot be easily extrapolated to females which is a limitation.

### The predictive values of memory contextualization for PTSD-like behavior

We hypothesized that insufficient contextualization of neutral trauma-related information, possibly due to a blunted HPA-axis response and hence impaired development of slow corticosteroid actions, is a predisposing trait for PTSD. This hypothesis is supported by indirect evidence which shows that cortisol administration after trauma reduced vulnerability to PTSD (Kothgassner et al., 2021; Matar et al., 2013; Schelling et al., 2004). We only partly confirmed this notion.

As mentioned, pre-trauma MC was predictive for PTSD susceptibility in the NS-group but not in the S-group. We cannot exclude that the difference in the incidence of PTSD-like behaviour meeting the cut-off criteria in the two experimental groups might have contributed to this observation. For example, the NS group included more animals with affected behaviour on *none* or *all* PTSD measures (than the S group), leading to a more diverse sample. While continuous values were used in the analyses, an alternative explanation for the observed group difference could be that MC is especially predictive for PTSD-like behaviour in samples with larger inter-individual variation. Although the observed group difference is contrary to the expectation based on earlier human studies, it would simplify testing individuals prior to trauma exposure for contextual memory ability–provided this finding can be translated to humans. Possibly, datasets are available in which contextual memory prior to trauma exposure was tested in humans under non-stressful conditions, to support the insights obtained in our current study. If MC had proved to be a stable trait, this might have further eased the applicability of such measurements to understand the relation with PTSD vulnerability, because in that case one could also test neutral MC post-trauma exposure. However, at least in animals, we found no support for stability of neutral MC in the face of trauma. Rather, the results suggest that trauma exposure may affect neutral MC performance.

In addition to the predefined hypotheses, the model also provided evidence for other significant correlations, most of which are in line with existing literature. Thus, pre-trauma trait anxiety was found to be positively related to PTSD-like behavior, as observed in humans (Kampman et al., 2017; Kok et al., 2016). Furthermore, peri-trauma freezing showed a positive relation to fear sensitization and association. One might speculate these relations representing freezing towards unconditioned and conditioned threat, respectively, reflect different, independent, neurobiological processes (i.e. innate fear or anxiety vs associative learning) (Hagenaars et al., 2014; Siegmund and Wotjak, 2007). Finally, PTSD-like behavior was positively related to fear association. This aligns with the increased contextual anxiety observed in PTSD-patients (Lissek and van Meurs, 2015).

While the fact that predictive value of neutral MC was particularly apparent in the NS-group already hints at a limited role for HPA-axis responsivity in the mediation of this effect, further testing also did not support a specific role of the HPA-axis. In fact, none of the significant correlations that we observed were mediated by HPA-axis variables. While recent literature indicates that blunted HPA-axis responsivity, likely due to increased negative feedback via glucocorticoid receptors, is a vulnerability factor for PTSD development after trauma exposure (Szeszko et al., 2018; Turner et al., 2020), this relation was not found in other studies or observed only in interaction with exposure to life adversity (Dunlop and Wong, 2019; Morris et al., 2016). This highlights the complex role of HPA-axis alterations in PTSD (Olff and van Zuiden, 2017; Schumacher et al., 2019). While we cannot exclude that our animal model did not capture this feature of PTSD–maybe related to less between-subject variability than one may expect in humans-, the data seems to indicate that HPA-axis abnormalities may add to PTSD-vulnerability independently from the currently investigated cognitive factor. This would align with recent models that identified problems with associative learning of neutral and trauma-related learning-due to impaired hippocampal function-as pre-trauma risk factor for PTSD (Alexander et al., 2020; Lambert and McLaughlin, 2019).

In conclusion, we demonstrated in rats that pre-trauma contextualization of neutral information and under non-stressful conditions significantly predicts the vulnerability to develop PTSD-like symptoms following trauma exposure. Future prospective studies in humans are necessary to investigate whether these findings in rodents can be extended to the human situation. If so, neutral memory contextualization might be added to the already known vulnerability factors (Bomyea et al., 2012; DiGangi et al., 2013; Tortella-Feliu et al., 2019; Yehuda and LeDoux, 2007), allowing to identify individuals at risk to develop PTSD, especially those having a higher likelihood to be exposed to trauma, such as soldiers deployed on a mission.

## Acknowledgements

We are thankful to Marijn Vellinga and Kate Liang for their vital help with the experimental set-up and pilot experiments, and to Sophie Heijnen and other colleagues for their crucial help with data collection. We thank the animal caretakers of our department for their assistance during experimental procedures, and Valeria Bonapersona, Jelle Knop and other colleagues for valuable discussions on the analysis. This work was supported by the Graduate program of the Netherlands Organization of Scientific Research [NWO grant number 022.003.003]; the Consortium on Individual Development (CID), which is funded through the Gravitation program of the Dutch Ministry of Education, Culture, and Science and the Netherlands Organization for Scientific Research [NWO grant number 024.001.003]; and the Dutch Ministry of Defense. The sponsors were not involved in the presented research.

## Author contributions

**Milou S.C. Sep**: Conceptualization, Methodology, Validation, Formal analysis, Investigation, Data Curation, Writing - Original Draft, Visualization, Project administration, Funding acquisition; **R. Angela. Sarabdjitsingh**: Conceptualization, Methodology, Validation, Resources, Writing - Review & Editing, Supervision, Project administration; **Elbert Geuze**: Conceptualization, Methodology, Resources, Writing - Review & Editing, Supervision, Funding acquisition; **Marian Joёls**: Conceptualization, Methodology, Validation, Resources, Writing - Original Draft, Supervision, Project administration, Funding acquisition

