## Appendix for "Pre-trauma memory contextualization as predictor for PTSD-like behavior in male rats"

### SUPPLEMENTAL INFORMATION (APPENDIX)

### A. Supplementary Methods

#### A1. Resident-intruder paradigm

Sprague Dawley (SD) rats were individually placed in the well-soiled home cage (Type IV Macrolon cage in use for 10 days) of an unfamiliar Long-Evans (LE) rat for 10 minutes. To ensure that this intervention is stressful but not traumatic (Calfa et al., 2006; Razzoli et al., 2009), the SD and LE rat were separated by a plexiglass plate with holes during exposure, to allow visual, auditory and olfactory, but not tactile, contact (Blanchard et al., 2001). LE rats were selected as residents based on previous research (Finnell et al., 2017; Reyes et al., 2015). Note, each experimental rat (in the S-group) encountered a different resident before and after trauma.

#### A2. Predator scent stress

Earlier studies used well-soiled -used for two days- cat litter in this model (e.g. refs (Cohen et al., 2020; Danan et al., 2018)). To minimize infection risk, we mixed sterile cat urine of domestic cats (obtained via bladder punctures and collected at the Veterinary Microbiological Diagnostic Centre, Faculty of Veterinary Medicine, Utrecht University) with sandbox sand (local hardware store) that was autoclaved in the research facility. Per batch (n=8), two closed transparent exposure cages (±25x25x40 cm) were prepared with 1200 g sand. To model exposure to 'a nearby cat in its territory', the sand was mixed with 1) 18 ml cat urine one day before exposure (left to air dray 24h), and 2) 18ml cat urine within 30 min before exposure. The total amount of urine per exposure box corresponded to a cat's average 2-day urine production per standard litter box filling (±55x40cm) (Pelligand et al., 2011; Zanghi et al., 2018).

#### A3. Elevated Plus Maze

The EPM is a, widely used, plus-shaped platform of grey plastic (60 cm above ground, arms: 50 cm length x 10 cm width) with two opposite open (light: 50 lux) and two opposite closed arms surrounded by 40 cm high opaque walls on three sides; 10 lux) (Walf and Frye, 2007). Rats were individually placed in the center of the maze, facing the north open arm, and allowed to freely explore the maze for 5 minutes. Each trial was videotaped, and movements were traced using behavioral tracking system EthoVision® XT 11.5 (Noldus Information Technology BV, Wageningen, The Netherlands). The EPM was cleaned with 70% ethanol after each animal. Time spent per zone and the number of arm entries (i.e. four-paws in arm) were used to calculate an anxiety index that integrates these two behavioral EPM parameters (Eq.(A.1), Appendix A7) (Cohen et al., 2018)).

#### A4. Acoustic Startle Responses

After the EPM, two animals at a time (cage mates) stayed 1.5 hours in their home cage before their Acoustic Startle Responses (ASR) were measured in two ventilated soundproof startle chambers (SR-LAB system, San Diego Instruments, San Diego, CA). To minimize signal disturbances by vibrations and noise, the chambers were placed on a stable weighing table and throughout the experiment a constant background sound was provided by a white noise generator (sound-level calibrated at 68db). SD rats were individually placed in a plexiglass cylinder resting on a vibration-sensitive platform (i.e. *Large animal enclosure* of SR-LAB system: 20.32 L x 8.89 I.D. cm) inside the startle chamber. A piezoelectric accelerometer below the platform detected (startle) movements inside the cylinder. To ensure consistent measurement sensitivity, the accelerometer was routinely calibrated to 200 mV +/- 5 mV for 25 trials using the SR-lab calibration unit. After a 5-minute acclimatization period, 30 acoustic startle stimuli (120 dB white noise; 40 ms) were delivered in six blocks (jittered inter-trial-interval 30-45 s) (Cohen et al., 2018). To ensure consistent presentation, sound levels within each chamber were routinely measured with a sound level meter. The startle amplitude was defined as the average response in the 100 ms response window following stimuli onset (Cohen et al., 2004). The cylinders were cleaned with soap and water after each animal. Mean startle amplitude (the 30 trial average) and percent habituation (Eq.(A.2), Appendix A7) were calculated (Cohen et al., 2018).

#### A5. Object-in-context task

##### A5.1 OIC set-up

In each OIC version, different pairs of objects and contexts were used to prevent learning effects (combinations counterbalanced across animals; https://osf.io/ztnva/). The two used object combinations -selected for equal preference in the pilot experiment- were 1) big ceramic mug (MUG) + glass laboratory staining dish with light bulb (GLASS) and 2) plastic cup (CUP) + Coca-Cola can (COLA). The selected objects had approximately equal size (about 7x12 cm), but differed in height, shape, and texture. Objects were secured to the context floor with Velcro tape (Roozendaal et al., 2008). The two used context combinations were 1) transparent plastic square boxes (69x46x39 cm) in OIC-square and 2) opaque plastic black round boxes (76x34 cm I.DxH) in OIC-round. To create distinct contexts, vertical lines of black tape were applied to one of the square boxes and a checkered pattern of squared white tape was applied to one of the round boxes. The floor of all contexts was covered with black painted standard Aspen Woodchips bedding (to create sufficient contrast between animal and box for automated tracking). One week prior to testing, the SD rats were habituated to this black bedding in their home cage.

##### A5.2 OIC procedures

Each version (i.e. OIC-round and OIC-square) consisted of 3 phases on 4 consecutive days. Note, in each phase, animals were placed in the context-boxes facing the same wall. On the first day (Habituation 1), SD rats were allowed to freely explore each version-specific context in 10-minute trials. This procedure was repeated on the next day (Habituation 2). On the third day (Training), each context contained a unique set of two identical, diagonally placed, objects (i.e. context X: objects A + A, context Y: objects B + B). The SD rats explored both object-context combinations in 10-minute trials. The SD rats in the S-group performed the OIC training phase 2.5h after the end of RI. The SD rats in the NS-group performed this phase without social stress. On the final day (Test), the SD rats were re-exposed to one of the contexts containing two unique objects (one from each training set, e.g. context X: objects A + B). The object that was encountered in the test context during the training phase is *in context* (object A), the other object is *out context* (object B) in the test phase. Rats explored the object-context combination in a 5-minute trial. In all phases, the context walls and -if applicable- objects were cleaned with 70% ethanol and -after removal of droppings- the bedding was mixed in between trials. Rats briefly returned to their home cage during this period. All animals had never been exposed to the objects and contexts prior to testing.

##### A5.3 OIC measures

For all phases, trials were recorded with an overhead video camera and movements were traced using behavioral tracking system EthoVision® XT 11.5 (Noldus Information Technology BV, Wageningen,

The Netherlands). Accurate identification of the nose-tail axis was manually checked via the track editor feature, if required manual nose-tail swaps were performed. Object exploration was defined as nose in proximity to the object (<2cm) or physical contact with the object via the rat's forepaws or snout (Bukhari et al., 2018; Czerniawski et al., 2015). Time spent per "*object zone"* was calculated in EthoVision and the absolute discrimination ratio (i.e. absolute MC index) was calculated (Eq. (A.3), Appendix A7): the absolute difference between time on out-context (novel) object and time on in-context (familiar) object, divided by the total time on objects.

#### A6. Pilot experiments to validate the experimental setup

**Table A1. Overview pilot experiments for experimental setup validation**

| **In text reference** | **Aim** | **Set-up** | **Animals** (42 SD and 4LE in total) | **Results** | **Summary Result** |
|---|---|---|---|---|---|
| Pilot I | Validation of blood corticosterone increase by RI and PSS procedures. Blood collected via tail cut, 30 min after stressor onset. (Each SD animal first exposed to RI and then to PSS) | RI, PSS | 6 SD & 2LE | Figure A1. | See Inconclusive results due to high corticosterone levels at baseline. Follow-up: Pilot IV, measurements in trunk blood |
| Pilot II | Test-retest reliability EPM and ASR with 18-day delay | EPM, ASR | 6 SD | Figure A2. and A3. | See Sufficient test-retest reliability. |
| Pilot III | Test-retest reliability OIC with 18-day delay | OIC | 6 SD (+ reused 6SD from pilot II) | Figure A4. | Good test-retest reliability. Follow-up: Pilot IV, validate the large individual variation in SD rats, and select the appropriate DR formula. |
| Pilot IV | Follow-up Pilot I and III. All animals (n=24 SD) performed the OIC once. One day later the animals were divided in 4 groups to measure corticosterone in trunk blood (after decapitation):<br>• At baseline (n=6). Decapitation from home cage.<br>• 45 minutes after onset RI (duration RI 10 minutes, n=6)<br>• 45 minutes after PSS onset (duration PSS 10 minutes, n=6<br>• 45 minutes after onset novelty stress (duration 30 minutes, n=6) | OIC, RI, PSS | (4x6=) 24 SD & 2 LE | Figure A5 | Large variation in OIC performance is consistently observed in SD rats. and the appropriate DR formula is selected.<br><br>Trunk blood samples showed that all stressors increased corticosterone levels. |

SD = male Sprague Dawley rats; LE = male Long Evans rats. Note the LE’s used during the pilot experiments were reused during the actual experiments.

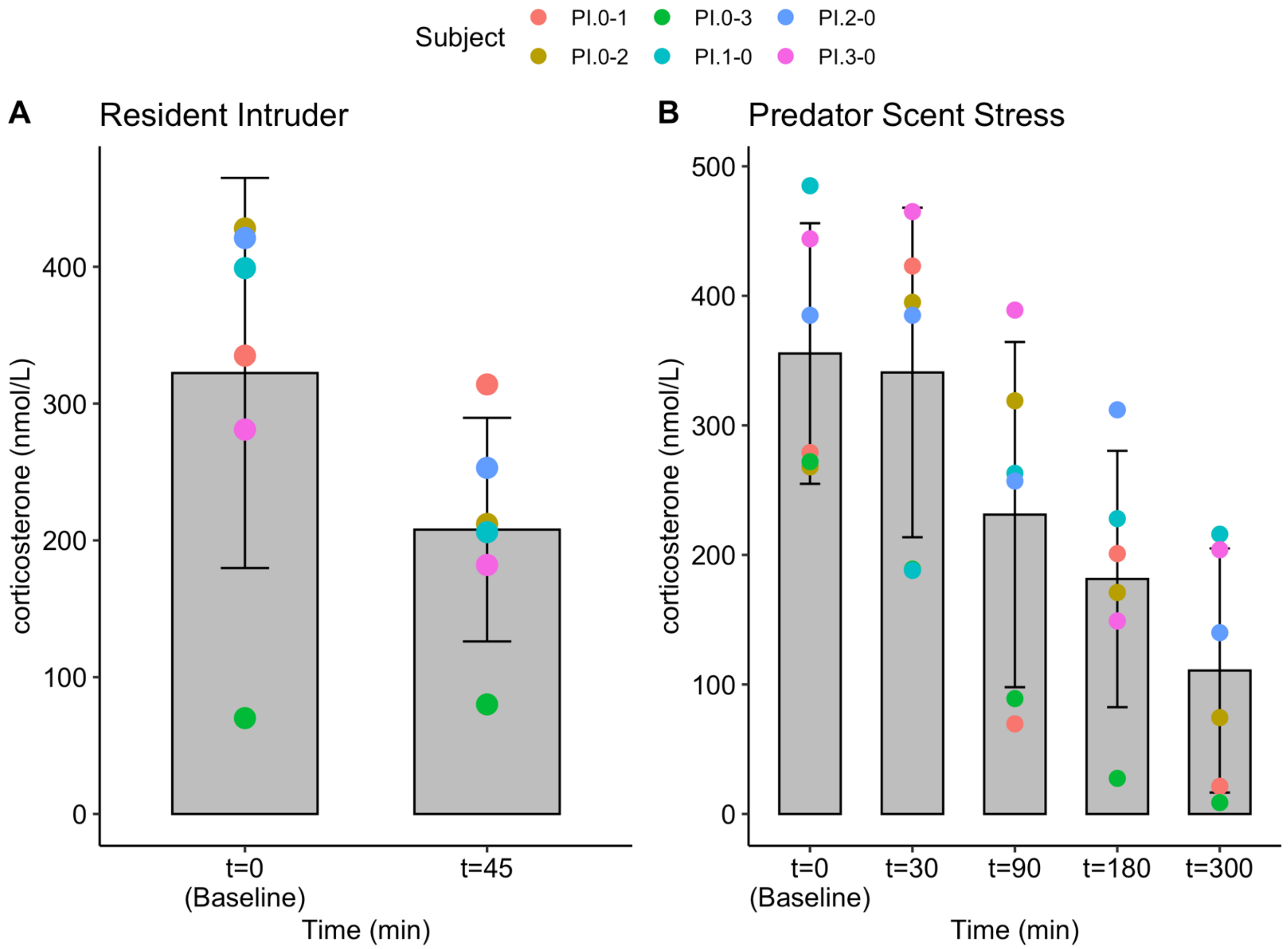

**Figure A1. Pilot I Serum corticosterone levels following Resident Intruder Stress and Predator Scent Stress.** Means and 95% CI. Blood samples were collected via tail cuts before and after RI (A) and PSS (B). Both graphs are inconclusive, due to high corticosterone values in the baseline groups. A new pilot (IV) was performed to evaluate corticosterone levels in trunk blood following RI and PSS.

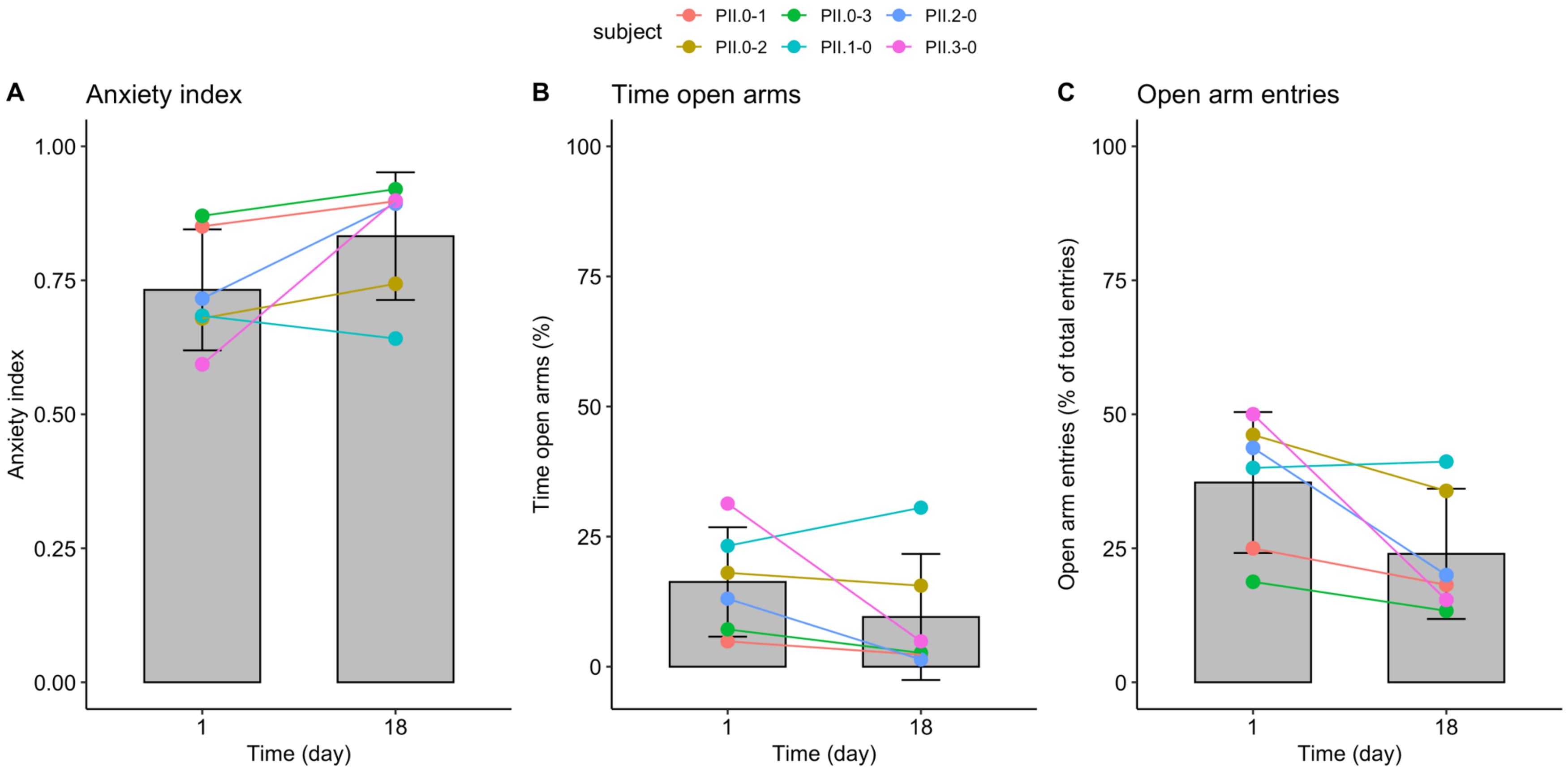

**Figure A2. Pilot II test-retest reliability of Elevated Plus Maze.** Means and 95% CI, dots represent individual measures. Paired t-test's did not reveal significant differences between day 1 and 18 on the anxiety index (A: $t(5)=-2.003$, $p=0.102$), time in open arms (B: $t(5)=1.443$, $p=0.209$) and open arm entries (C: $t(5)=2.45$, $p=0.056$). Based on these findings, we concluded that the EPM has good test-retest reliability. Anxiety index: Eq. (A.1), Appendix A7.

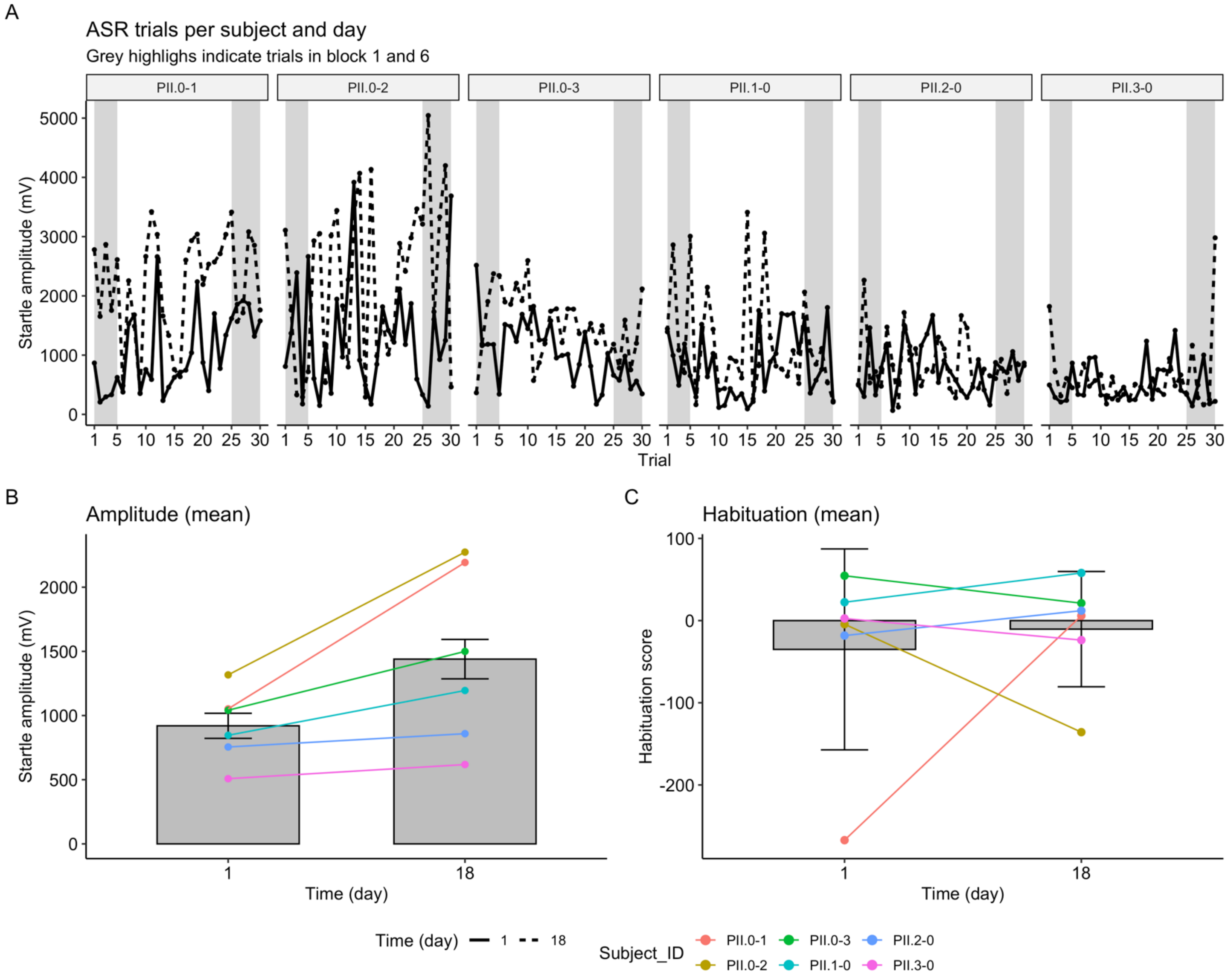

**Figure A3. Pilot II test-retest reliability of Acoustic Startle Responses.** For each animal, ASR amplitudes per trial are shown for both experimental days (A). Trials in block 1 and 6 were used to calculate the habituation score (Eq.(A.2), Appendix A7). Paired t-test's reveal a significant difference between day 1 and 18 on amplitude (B: $t(5)=-2.920$, $p=0.033$), but not on habituation index (C: $t(5)=-0.445$, $p=0.6752$). B and C: means and 95% CI, dots represent individual measures. Plot A and B indicate that only two animals showed increased ASR amplitudes on day 18, while the amplitude remained constant between day 1 and 18 for the other 4 animals. Based on these findings, we concluded that the ASR has sufficient test-retest reliability.

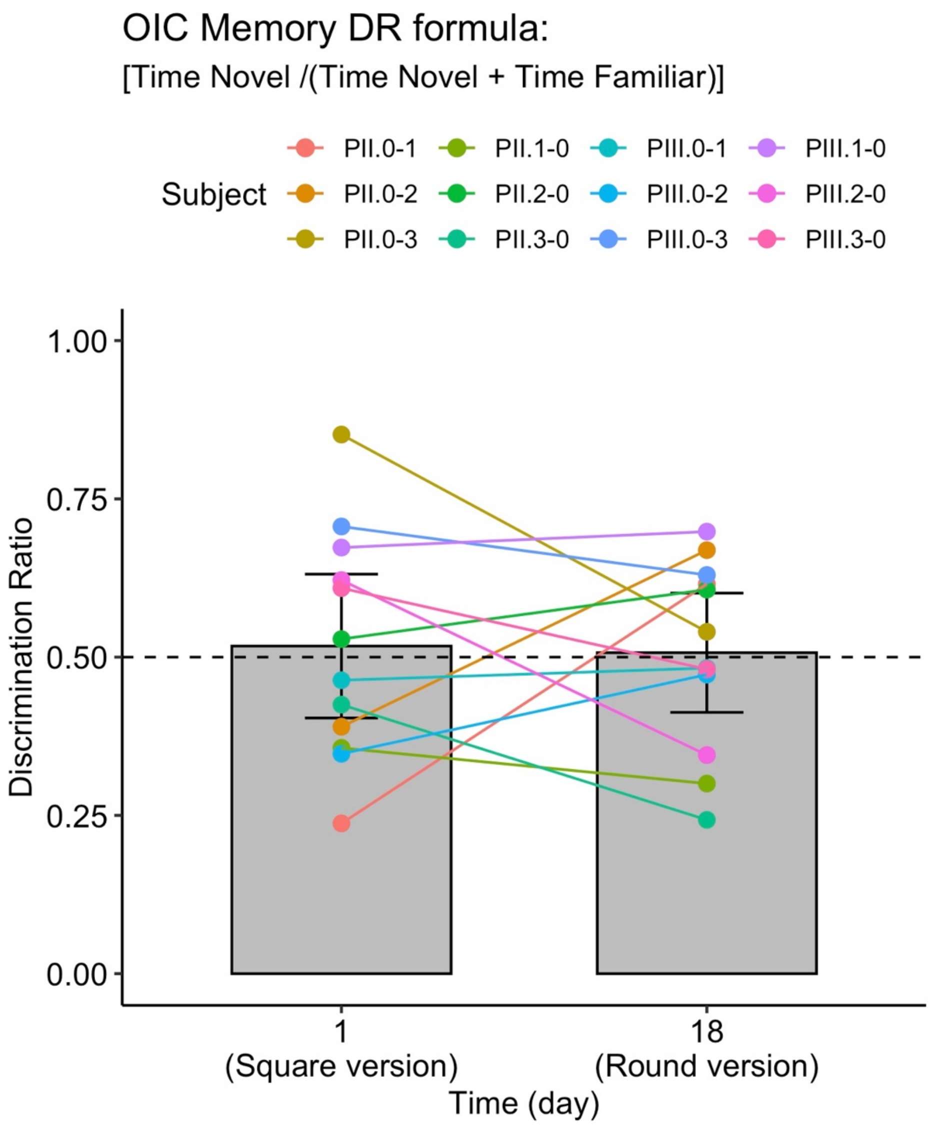

**Figure A4. Pilot III test-retest reliability of the Object-in-context task.** Means and 95% CI, dots represent individual measures. Subjects with prefix PIII were test-naïve, subjects with prefix PII had previously performed pilot II. Paired t-test revealed no significant difference in DR between day 1 and day 18 (t(11)=0.176, p = 0.864). A one sample t-test showed that the DR in the total sample was not different from 0.5 (t(23)=0. 372, p=0.713). Based on these results we concluded that test-retest reliability of the OIC versions was good. A follow-up pilot (IV) was performed to validate that large variation and average group performance around 0.5 a consistent observation in SD rats.

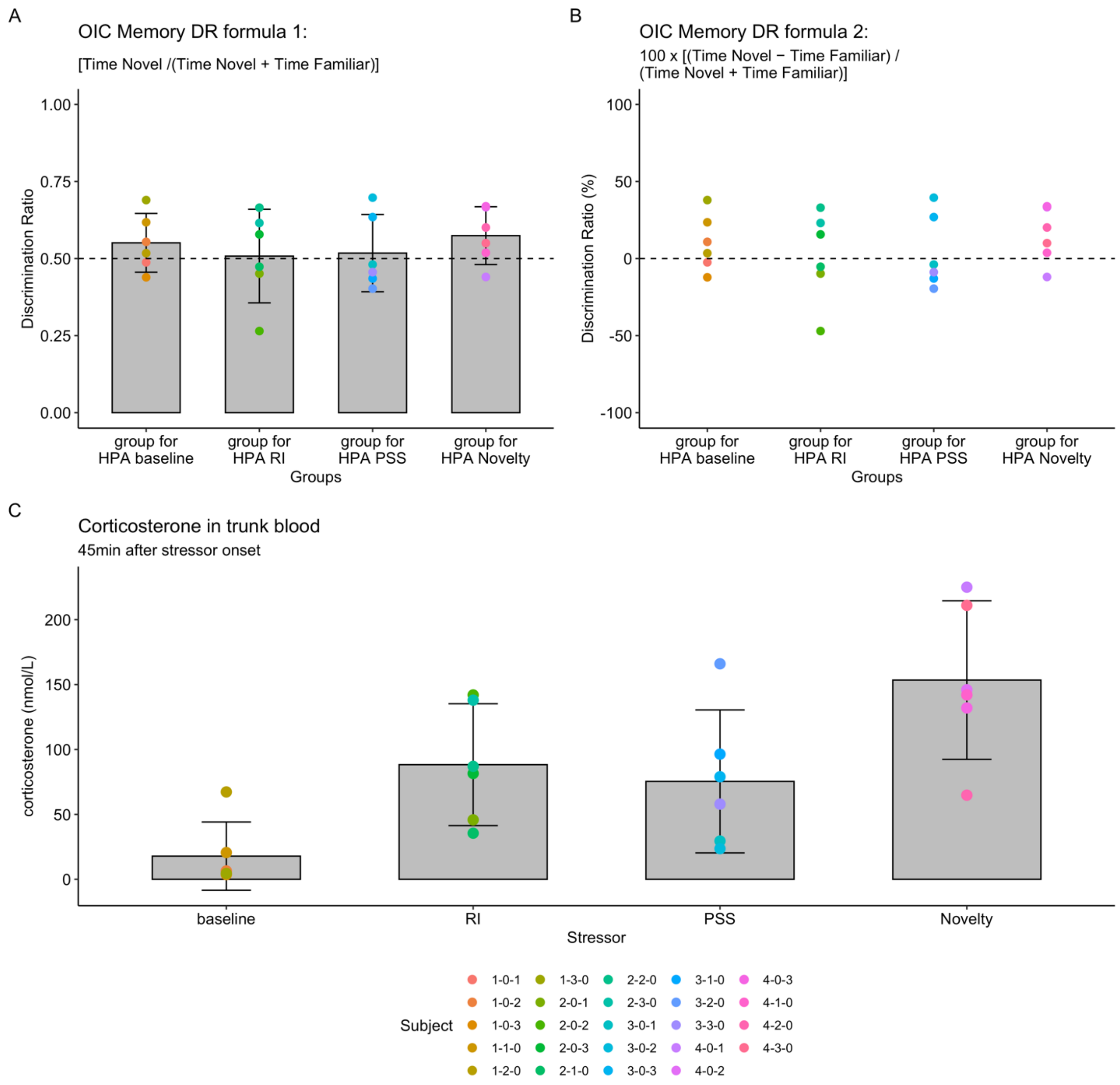

**Figure A5. Pilot IV: validation Object-in-Context task variation and stressor induced corticosterone responses.** All animals in this pilot were first exposed to the OIC and then divided in 4 groups for corticosterone measures. Dots represent individual measures. A and C: means and 95% CI. The OIC results in this pilot aligned with the observation in pilot III: individual variation is substantial and average group performance was around 0.5 - i.e. the DR in the total sample was not different form 0.5 (t(23)=1.697, p=0.103)- (A). Two formulas were used to calculate the DR ratio in this pilot: centered around 0.5 (A) and centered around 0 (B). The consistently observed variation in SD rats suggests that the (absolute value of the) DR formula that centers around 0 (B) is more appropriate to capture performance in SD rats. Using this formula, performance around 0 is interpreted as 'no contextual memory', while DR's that differ from 0 (regardless of the sign) reflect the formation of context-dependent memory (Sep et al., 2021). Pairwise one-sided t-tests (with Benjamini-Hochberg correction for multiple testing, and pooled SD due to limited sample size) showed that the corticosterone levels in trunk blood were reliable increased by the RI (p=0.017), PSS (p=0.028) and Novelty (p<0.001) stress procedures (C).

#### A7. EQUATIONS

Eq. (A.1)

$$Anxiety\ index = 1 - \left( \frac{\left( \frac{time\ spent\ in\ the\ open\ arms}{total\ time\ in\ the\ maze} + \frac{number\ of\ entries\ to\ the\ open\ arms}{total\ entries\ open\ and\ closed\ arms} \right)}{2} \right)$$

Eq. (A.2)

$$Habituation\ index = 100 * \frac{(average\ startle\ amplitude\ block\ 1 - average\ startle\ amplitude\ block\ 6)}{average\ startle\ amplitude\ block\ 1}$$

Eq. (A.3)

$$absolute\ MC\ index = \left( \frac{abs(time\ out\ context - time\ in\ context)}{(time\ out\ context + time\ in\ context)} \right)$$

#### A8. Hypothesized direct and indirect relations

**Table A2. Hypothesized direct and indirect relations**

| **Research questions and hypothesis** | **Modelled paths and estimated parameters** | **References (if applicable)** |
|---|---|---|
| **A. Hypothesized relations to answer the primary research questions** | | |
| Is pre-trauma neutral MC a risk factor for PTSD? | • PTSD-like behavior ~ (a1,a2) * Pre-trauma MC | |
| Is neutral MC a stable trait? | • Post-trauma MC ~ (b1,b2) * Pre-trauma MC | |
| Is the relation between pre- and post-trauma neutral MC mediated by PTSD? | • Post-trauma MC ~ (c1,c2) * PTSD-like behavior<br>*Mediation 1: Pre-trauma MC -> PTSD-like -> Post-trauma MC*<br>• *group 1 := a1 * c1*<br>• *group 2 := a2 * c2* | |
| Is pre-trauma neutral MC related to post-trauma fear sensitization / association? | • Fear association ~ (d1,d2) * Pre-trauma MC<br>• Fear sensitization ~ (e1,e2) * Pre-trauma MC | |
| Is the relation between PTSD and post-trauma fear sensitization and association mediated by post-trauma neutral MC? | • Fear association ~ c(f,f) * PTSD-like behavior<br>• Fear sensitization ~ c(g,g)* PTSD-like behavior<br>• Fear association ~ c(d11,d22) * post-trauma MC<br>• Fear sensitization ~ c(e11,e22) * post-trauma MC<br>*Mediation 2: PTSD-like -> post-trauma MC -> fear association*<br>• *group 1 := c1 * d11*<br>• *group 2 := c2 * d22*<br>*Mediation 3: PTSD-like -> post-trauma MC -> fear sensitization*<br>• *group 1 := c1 * e11*<br>• *group 2 := c2 * e22* | |
| Is HPA-axis reactivity during trauma related to subsequent PTSD susceptibility? | PTSD-like behavior ~ (h,h) * Peri-trauma corticosterone | |
| Is the relation between pre-trauma neutral MC and PTSD mediated by HPA-axis reactivity during trauma? | • Peri-trauma corticosterone ~ (i1,i2) * Pre-trauma MC<br>*Mediation 4: Pre-trauma MC -> corticosterone -> PTSD-like* | |

| | | |
|---|---|---|
| | • *group 1 := i1 * h*<br>• *group 2 := i2 * h* | |
| **B Relations based on previous findings or experimental design** | | |
| B1. Fear during trauma | | (Hagenaars et al., 2014; Siegmund and Wotjak, 2007) |
| 1. Freezing during trauma and freezing during re-exposure | • Fear sensitization ~ (m,m) * peri-trauma freezing<br>• Fear association ~ (n,n) * peri-trauma freezing | |
| 2. Freezing during trauma and PTSD susceptibility | • PTSD-like behaviour ~ (o,o) * peri-trauma freezing | |
| B2. HPA & learning and memory | | (Sep et al., 2019a, 2019c) |
| HPA-reactivity can have time-dependent effects on memory contextualization, and might mediate the relation between pre-trauma neutral MC and trauma MC. | • Fear sensitization ~ (x,x) * Peri-trauma corticosterone<br>• Fear association ~ (y,y) * Peri-trauma corticosterone<br><br>*Mediation 5: Pre-trauma MC -> corticosterone -> Fear association*<br><br>• *group 1 := i1 * y*<br>• *group 2 := i2 * y*<br><br>*Mediation 6: Pre-trauma MC -> corticosterone -> Fear sensitization*<br><br>• *group 1 := i1 * x*<br>• *group 2 := i2 * x* | |
| B3. Trait Anxiety (TA): | | |
| 1. *Fearful MC (following acute stress) can be influenced by the interaction between TA and HPA-reactivity* | • Pre-trauma MC ~ (aa1,aa2) * TA<br><br>*Mediation 7: TA -> corticosterone -> Fear association*<br><br>• *group 1+2 := l * y*<br><br>*Mediation 8: TA -> corticosterone -> Fear sensitization*<br><br>• *group 1+2 := l * x* | (Sep et al., 2020) |
| 2. *TA is associated with an increased risk to develop PTSD* | • PTSD-like behavior ~ (j,j) * TA | (Kampman et al., 2017; Kok et al., 2016) |

| | | |
|---|---|---|
| 3. *HPA-axis reactivity and fear expression during trauma can be influenced by TA* | • Peri-trauma freezing ~ (k,k) * TA<br>• Peri-trauma corticosterone ~ (l,l) * TA | (Weger and Sandi, 2018); Fear expression (Craske et al., 2009); HPA-reactivity (Hauner et al., 2008) |
| 4. *TA can influence fear generalization/sensitization* | • Fear sensitization ~ (z,z) * TA | (Sep et al., 2019b) |
| **C. Experimental design** | | |
| C1. Covariance between variables that are measured on the same experimental day | | Not applicable |
| 1. peri-trauma measures | • Peri-trauma freezing ~~ (p,p) * Peri-trauma corticosterone | |
| 2. post trauma MC indices (context-dependent memory and generalization) | • Fear sensitization ~~ (q,q) * Fear association | |
| C2. Batch as covariate<br><br>Note, two parameters were estimated for the effects of batch (indicated with bb and bbc), as correlation matrixes revealed a correlation of batch with Fear association and Fear sensitization, but not with other variables. | • TA ~ (bb,bb) * batch<br>• PTSD-like behaviour ~ (bb,bb) * batch<br>• Pre-trauma MC ~ (bb,bb) * batch<br>• Post-trauma MC ~ (bb,bb) * batch<br>• peri-trauma freezing ~ (bb,bb) * batch<br>• Peri-trauma corticosterone ~ (bb,bb) * batch<br><br>• Fear sensitization ~ (bbc,bbc) * batch<br>• Fear association ~ (bbc,bbc) * batch | Not applicable |
| *Note, letter-codes between brackets "()" in modelled paths column indicate estimated parameters for both experimental groups, separated by a comma. Same letter-codes indicate that estimated parameters were constrained to be equal between groups, different letter-codes indicate that the estimated parameters are estimated within experimental group. Operators in lavaan syntax* (Rosseel, 2012)*: ~ direct effects (regressions), := Indirect effects (mediation), ~~ (co)variance* | | |

### B. Supplementary Results

#### B1. Estimated direct and indirect paths

**Table B1. Estimated parameters of direct and indirect paths per experimental group**

| Group | Outcome | Indicator | Path Label | B | SE | Z | CI lower | CI upper | Beta | p-value | significance |
|---|---|---|---|---|---|---|---|---|---|---|---|
| **NS** | PTSD-like behavior | pre-trauma MC | a1 | -1,827 | 0,912 | -2,003 | -3,810 | -0,196 | -0,283 | 0,045 | * |
| **NS** | post-trauma MC | pre-trauma MC | b1 | -0,013 | 0,145 | -0,087 | -0,284 | 0,286 | -0,016 | 0,931 | |
| **NS** | post-trauma MC | PTSD-like behavior | c1 | 0,012 | 0,024 | 0,476 | -0,043 | 0,054 | 0,094 | 0,634 | |
| **NS** | fear association | pre-trauma MC | d1 | 0,237 | 0,098 | 2,425 | 0,052 | 0,438 | 0,351 | 0,015 | * |
| **NS** | fear sensitization | pre-trauma MC | e1 | -0,027 | 0,102 | -0,262 | -0,239 | 0,163 | -0,035 | 0,793 | |
| **NS** | fear association | PTSD-like behavior | f | 0,029 | 0,014 | 2,119 | 0,002 | 0,056 | 0,282 | 0,034 | * |
| **NS** | fear sensitization | PTSD-like behavior | g | 0,010 | 0,018 | 0,541 | -0,024 | 0,049 | 0,084 | 0,588 | |
| **NS** | fear association | post-trauma MC | d11 | -0,039 | 0,118 | -0,332 | -0,283 | 0,176 | -0,046 | 0,740 | |
| **NS** | fear sensitization | post-trauma MC | e11 | 0,050 | 0,134 | 0,373 | -0,218 | 0,308 | 0,052 | 0,709 | |
| **NS** | PTSD-like behavior | peri-trauma corticosterone | h | 0,640 | 1,789 | 0,358 | -2,543 | 4,380 | 0,035 | 0,720 | |
| **NS** | peri-trauma corticosterone | pre-trauma MC | i1 | -0,091 | 0,055 | -1,654 | -0,206 | 0,011 | -0,260 | 0,098 | + |
| **NS** | fear sensitization | peri-trauma corticosterone | x | 0,080 | 0,181 | 0,441 | -0,318 | 0,406 | 0,036 | 0,659 | |
| **NS** | fear association | peri-trauma corticosterone | y | 0,143 | 0,242 | 0,590 | -0,280 | 0,695 | 0,074 | 0,555 | |
| **NS** | pre-trauma MC | trait anxiety | aa1 | -0,033 | 0,037 | -0,898 | -0,096 | 0,048 | -0,158 | 0,369 | |
| **NS** | PTSD-like behavior | trait anxiety | j | 0,637 | 0,117 | 5,467 | 0,415 | 0,880 | 0,469 | 0,000 | *** |

| | | | | | | | | | | | |
|---|---|---|---|---|---|---|---|---|---|---|---|
| **NS** | peri-trauma freezing | trait anxiety | k | 0,024 | 0,022 | 1,134 | -0,021 | 0,064 | 0,136 | 0,257 | |
| **NS** | peri-trauma corticosterone | trait anxiety | l | -0,007 | 0,009 | -0,869 | -0,026 | 0,008 | -0,102 | 0,385 | |
| **NS** | fear sensitization | trait anxiety | z | 0,006 | 0,018 | 0,308 | -0,039 | 0,036 | 0,035 | 0,758 | |
| **NS** | fear sensitization | peri-trauma freezing | m | 0,303 | 0,103 | 2,941 | 0,098 | 0,493 | 0,341 | 0,003 | ** |
| **NS** | fear association | peri-trauma freezing | n | 0,380 | 0,111 | 3,430 | 0,195 | 0,635 | 0,483 | 0,001 | *** |
| **NS** | PTSD-like behavior | peri-trauma freezing | o | -1,476 | 0,905 | -1,631 | -3,335 | 0,217 | -0,196 | 0,103 | |
| **S** | PTSD-like behavior | pre-trauma MC | a2 | 0,075 | 0,929 | 0,081 | -1,820 | 1,888 | 0,011 | 0,936 | |
| **S** | post-trauma MC | pre-trauma MC | b2 | 0,048 | 0,133 | 0,358 | -0,212 | 0,313 | 0,058 | 0,720 | |
| **S** | post-trauma MC | PTSD-like behavior | c2 | 0,006 | 0,031 | 0,178 | -0,056 | 0,063 | 0,045 | 0,859 | |
| **S** | fear association | pre-trauma MC | d2 | -0,120 | 0,147 | -0,814 | -0,453 | 0,128 | -0,132 | 0,415 | |
| **S** | fear sensitization | pre-trauma MC | e2 | -0,016 | 0,105 | -0,155 | -0,222 | 0,191 | -0,024 | 0,877 | |
| **S** | fear association | PTSD-like behavior | f | 0,029 | 0,014 | 2,119 | 0,002 | 0,056 | 0,215 | 0,034 | * |
| **S** | fear sensitization | PTSD-like behavior | g | 0,010 | 0,018 | 0,541 | -0,024 | 0,049 | 0,100 | 0,588 | |
| **S** | fear association | post-trauma MC | d22 | -0,154 | 0,160 | -0,963 | -0,477 | 0,159 | -0,139 | 0,336 | |
| **S** | fear sensitization | post-trauma MC | e22 | -0,003 | 0,160 | -0,021 | -0,306 | 0,319 | -0,004 | 0,983 | |
| **S** | PTSD-like behavior | peri-trauma corticosterone | h | 0,640 | 1,789 | 0,358 | -2,543 | 4,380 | 0,053 | 0,720 | |
| **S** | peri-trauma corticosterone | pre-trauma MC | i2 | 0,135 | 0,099 | 1,360 | -0,066 | 0,327 | 0,246 | 0,174 | |
| **S** | fear sensitization | peri-trauma corticosterone | x | 0,080 | 0,181 | 0,441 | -0,318 | 0,406 | 0,066 | 0,659 | |
| **S** | fear association | peri-trauma corticosterone | y | 0,143 | 0,242 | 0,590 | -0,280 | 0,695 | 0,086 | 0,555 | |
| **S** | pre-trauma MC | trait anxiety | aa2 | 0,006 | 0,036 | 0,167 | -0,064 | 0,077 | 0,038 | 0,868 | |

| | | | | | | | | | | | |
|---|---|---|---|---|---|---|---|---|---|---|---|
| **S** | PTSD-like behavior | trait anxiety | j | 0,637 | 0,117 | 5,467 | 0,415 | 0,880 | 0,609 | 0,000 | *** |
| **S** | peri-trauma freezing | trait anxiety | k | 0,024 | 0,022 | 1,134 | -0,021 | 0,064 | 0,171 | 0,257 | |
| **S** | peri-trauma corticosterone | trait anxiety | l | -0,007 | 0,009 | -0,869 | -0,026 | 0,008 | -0,086 | 0,385 | |
| **S** | fear sensitization | trait anxiety | z | 0,006 | 0,018 | 0,308 | -0,039 | 0,036 | 0,054 | 0,758 | |
| **S** | fear sensitization | peri-trauma freezing | m | 0,303 | 0,103 | 2,941 | 0,098 | 0,493 | 0,416 | 0,003 | ** |
| **S** | fear association | peri-trauma freezing | n | 0,380 | 0,111 | 3,430 | 0,195 | 0,635 | 0,380 | 0,001 | *** |
| **S** | PTSD-like behavior | peri-trauma freezing | o | -1,476 | 0,905 | -1,631 | -3,335 | 0,217 | -0,202 | 0,103 | |

| Group | Mediations | Path Label | B | SE | Z | CI lower | CI upper | Beta | p-value | significance |
|---|---|---|---|---|---|---|---|---|---|---|
| **NS** | Mediation 1: Pre-trauma MC -> PTSD-like -> Post-trauma MC | a1*c1 | -0,021 | 0,050 | -0,421 | -0,152 | 0,058 | -0,027 | 0,674 | |
| **S** | Mediation 1: Pre-trauma MC -> PTSD-like -> Post-trauma MC | a2*c2 | 0,000 | 0,029 | 0,014 | -0,063 | 0,063 | 0,001 | 0,989 | |
| **NS** | Mediation 2: PTSD-like -> post-trauma MC -> Fear association | c1*d11 | 0,000 | 0,003 | -0,136 | -0,011 | 0,004 | -0,004 | 0,892 | |
| **S** | Mediation 2: PTSD-like -> post-trauma MC -> Fear association | c2*d22 | -0,001 | 0,007 | -0,116 | -0,025 | 0,009 | -0,006 | 0,908 | |
| **NS** | Mediation 3: PTSD-like -> post-trauma MC -> Fear sensitization | c1*e11 | 0,001 | 0,004 | 0,147 | -0,004 | 0,013 | 0,005 | 0,883 | |
| **S** | Mediation 3: PTSD-like -> post-trauma MC -> Fear sensitization | c2*e22 | 0,000 | 0,005 | -0,003 | -0,013 | 0,010 | 0,000 | 0,997 | |
| **NS** | Mediation 4: Pre-trauma MC -> corticosterone -> PTSD-like | i1*h | -0,058 | 0,195 | -0,298 | -0,633 | 0,192 | -0,009 | 0,766 | |
| **S** | Mediation 4: Pre-trauma MC -> corticosterone -> PTSD-like | i2*h | 0,086 | 0,300 | 0,288 | -0,299 | 1,086 | 0,008 | 0,773 | |
| **NS** | Mediation 5: Pre-trauma MC -> corticosterone -> Fear association | i1*y | -0,013 | 0,024 | -0,541 | -0,085 | 0,019 | -0,019 | 0,588 | |
| **S** | Mediation 5: Pre-trauma MC -> corticosterone -> Fear association | i2*y | 0,019 | 0,041 | 0,471 | -0,026 | 0,165 | 0,018 | 0,637 | |

| | | | | | | | | | |
|---|---|---|---|---|---|---|---|---|---|
| **NS** | Mediation 6: Pre-trauma MC -> corticosterone -> Fear sensitization | i1*x | -0,007 | 0,019 | -0,379 | -0,057 | 0,023 | -0,009 | 0,705 |
| **S** | Mediation 6: Pre-trauma MC -> corticosterone -> Fear sensitization | i2*x | 0,011 | 0,031 | 0,348 | -0,031 | 0,104 | 0,009 | 0,728 |
| **NS + S** | Mediation 7: TA -> corticosterone -> Fear association | I*y | -0,001 | 0,003 | -0,370 | -0,012 | 0,002 | -0,007 | 0,712 |
| **NS + S** | Mediation 8: TA -> corticosterone -> Fear sensitization | I*x | -0,001 | 0,002 | -0,264 | -0,008 | 0,002 | -0,004 | 0,792 |

*B: unstandardized path-coefficients. SE: bootstrapped standard errors of B. Z: z-statistic = B divided by SE. CI lower and CI upper: bootstrapped bias-corrected and accelerated 95% Confidence Interval of B. p-value: p-value corresponding to Z. Beta: standardized path-coefficients. Significance indicators: p-value < .1 (+), p-value < .05 (*), p-value < .01 (**), p-value < .001 (***). Note batch was included in the model as covariate (i.e. relations with this variable were not of primary interest to answer the research question at hand), for brevity the estimated parameters for batch are not shown in this table (significant relations are shown in figure 5).*

#### B2. Follow-up analysis group-differences MC

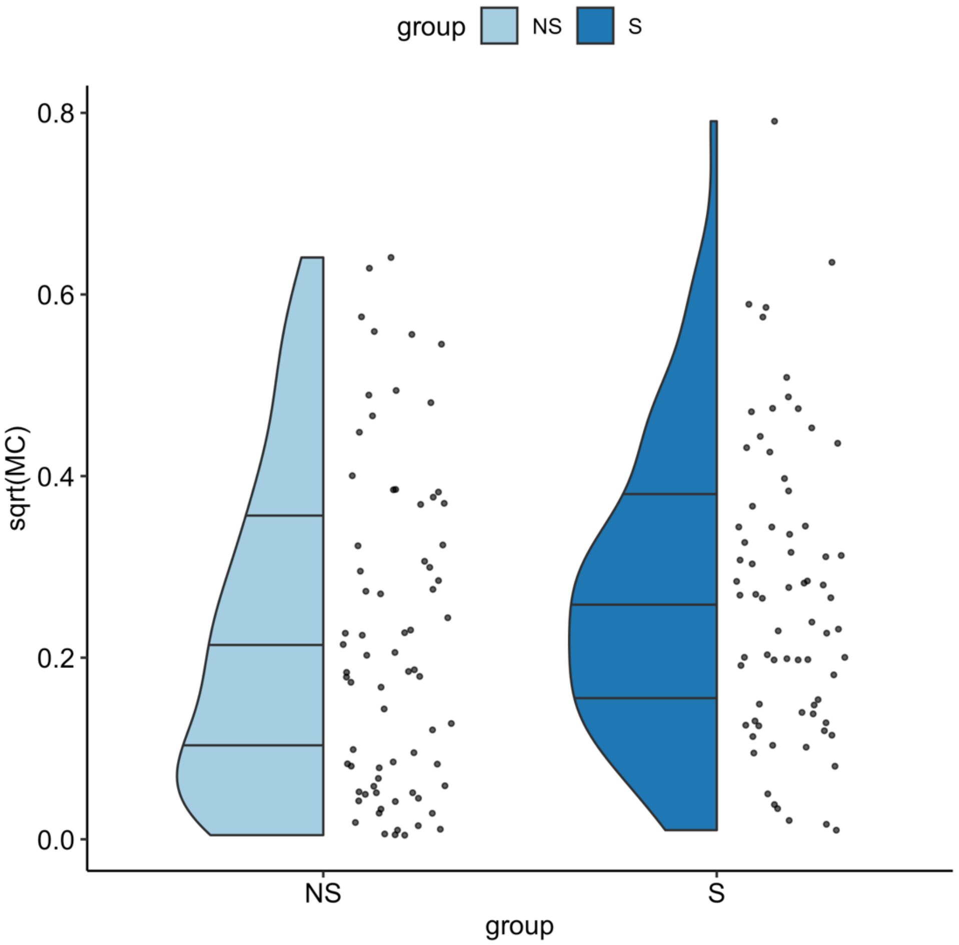

**Figure B1. Exploration of MC in NS- and S-groups.** Explorative comparison of memory contextualization (MC) in both experimental groups revealed that MC was better in the S-group compared to the NS-group (F(1)=4.916, p=0.028). Since path-analysis suggest that pre- and post-trauma MC measures represent independent variables, these measures were combined to increase the sample size per experimental group. Dots indicate individual datapoints.
